# Human Saliva Induces Acetoin-Dependent Metabolic Adaptation that Promotes Stress Tolerance in *Streptococcus mutans*

**DOI:** 10.64898/2026.10.27.762223

**Authors:** Jacob S. Harris, Alyssa N. Deever, Kyulim Lee, Justin R. Kaspar

## Abstract

Oral microbes are continually exposed to host-derived saliva, yet its effects on bacterial metabolism and stress adaptation remain incompletely defined. We investigated how saliva supplementation alters the physiology of the caries-associated bacterium *Streptococcus mutans* using transcriptomics (RNA-Seq), targeted metabolite measurements, specific gene deletions, and transposon sequencing (Tn-Seq). Growth in a chemically defined medium supplemented with pooled saliva altered expression of genes involved in carbohydrate transport, purine biosynthesis, and pyruvate-associated metabolism, including induction of the acetoin dehydrogenase operon. Saliva significantly increased extracellular acetoin accumulation across all *S. mutans* isolates tested, whereas little accumulation occurred in two commensal streptococcal species examined. Deletion of acetolactate synthase/decarboxylase (*alsSaldB*) abolished the saliva-associated increase, while deletion of individual acetoin dehydrogenase components (*adhC* /*lplA*) enhanced accumulation, supporting the predicted roles of genes in acetoin synthesis and catabolism. Increasing the available carbohydrate concentrations within the media further increased acetoin accumulation, alongside declining terminal culture pH that was significantly lower in cultures supplemented with saliva. Following growth with saliva, *S. mutans* exhibited approximately 10,000-fold greater survival during a hydrogen peroxide challenge than cells grown without saliva. Mutants within the acetoin pathway retained saliva-associated protection but showed approximately 100-fold lower survival than the wild type. Serial passage of a pooled transposon library identified previously unrecognized saliva-associated fitness determinants, including depletion of mutants carrying insertions in the acetoin dehydrogenase operon. Together, these findings identify acetoin metabolism as a contributor to saliva-associated metabolic remodeling and oxidative stress tolerance in *S. mutans* and establish saliva as a critical modulator of *S. mutans* physiology.

## INTRODUCTION

Oral microbes localized to the hard, non-shedding surfaces of the tooth inhabit supragingival biofilms that are continually exposed to human saliva. Host-derived saliva supplies glycoproteins, peptides, amino acids, and other substrates that can support microbial metabolism in a resource-limited environment, while its antimicrobial activities and effects on attachment can simultaneously constrain bacterial growth and colonization (1–4). These opposing influences make saliva an important determinant of microbial persistence and community behavior in the oral cavity. Studies of individual salivary components have previously demonstrated that oral bacteria actively respond to this environment: recognition of the mucin MUC5B alters adhesin expression and pathways associated with glycan utilization in *Streptococcus gordonii*, while salivary mucins and their glycans can promote bacterial coexistence and maintain oral community diversity (5–7). However, the physiological consequences of exposing oral bacteria to the combined components of saliva remain poorly defined, and conventional microbiological media do not fully reproduce these host-derived influences.

*Streptococcus mutans* is a prominent caries-associated bacterium whose ability to ferment dietary carbohydrates and tolerate acidic conditions favors its persistence as supragingival biofilms acidify (8). Repeated carbohydrate exposure from host feeding and the resulting decrease in biofilm pH can select for acidogenic and acid-tolerant organisms, promoting conditions that drive tooth enamel demineralization (9–11). The outputs of pyruvate metabolism directly connect carbohydrate utilization to this environmental acidification that can destabilize the microbiome. Depending on nutrient availability and oxygen exposure, *S. mutans* can direct pyruvate toward lactate or mixed-acid fermentation products, including formate, acetate, and ethanol (12–15). Changes in carbohydrate transport and pyruvate partitioning can also alter acid production and competitive fitness in mixed-species settings (16). Consequently, evaluating *S. mutan*s metabolism in the presence of saliva may reveal physiological states and fitness impacts that have not been apparent during growth in standard laboratory media.

Successful persistence within this competitive environment also requires *S. mutans* to withstand oxidative stressors, including hydrogen peroxide produced by neighboring oral streptococci (17, 18). Although *S. mutans* lacks catalase, its antioxidant defenses include peroxide-reducing enzymes such as AhpCF and Tpx, as well as iron sequestration by Dpr, which limits the formation of damaging hydroxyl radicals (19, 20). These defenses are regulated in part by the Spx proteins: SpxA1 serves as the major activator of oxidative stress genes, whereas SpxA2 contributes to oxidative and cell envelope stress tolerance (19, 21). The Spx-regulated response also encompasses changes in pyruvate metabolism, including induction of acetoin pathway genes and increased acetoin accumulation in an SpxA1-dependent manner (22). Acetoin is a nonacidic fermentation product formed from the condensation of pyruvate through acetolactate, and the Adh enzyme complex is implicated in its catabolism (22, 23). The AdhC subunit also interacts with SpxA2, suggesting a further connection between acetoin-associated enzymes and stress regulation, although the physiological significance of this interaction remains unresolved (23).

Our previous work demonstrated that saliva supplementation substantially alters the physiology of several oral streptococci, including *S. mutans* (24). Addition of saliva shortened the *S. mutans* lag phase and doubling time and modulated expression of genes involved in carbohydrate uptake and biofilm formation, including *gtfB* and *gtfC*, which encode glucan-producing glucosyltransferases. However, the most striking phenotype documented was the enhanced competitiveness of *S. mutans* in saliva against commensal streptococci; *S. mutans* displayed a 76-fold increase in competitive index against *Streptococcus oralis* and a 21-fold increase against *Streptococcus sanguinis* (24). In this study, we sought to identify the genetic determinants that contribute to enhanced fitness under these conditions. Our findings reveal that saliva alters metabolic and transcriptional activity in *S. mutans*, including changes in pyruvate metabolism and increased acetoin accumulation. Consistent with previous reports, these results support a contribution of acetoin metabolism to oxidative stress tolerance and demonstrate that growth with saliva markedly enhances tolerance to hydrogen peroxide. Together, these findings provide insight into how host-derived environmental factors shape *S. mutans* physiology and identify metabolic pathways that contribute to its fitness under saliva-supplemented conditions.

## RESULTS

### Saliva supplementation alters metabolic gene expression in Streptococcus mutans

Our previous work demonstrated that supplementation with pooled human saliva alters the transcriptional profiles of oral streptococci, including *S. mutans* (24). These experiments were performed in tryptone–yeast extract (TY) broth, a complex medium that has an incompletely defined nutrient composition. We therefore examined *S. mutans* in a chemically defined medium (CDM) (25, 26) to characterize saliva-associated transcriptional changes against a defined nutritional background and determine whether responses previously observed in TY were retained. RNA was isolated from mid-logarithmic phase cultures grown in CDM without saliva (-HS) or supplemented with 50% (v/v) pooled human saliva (+HS), and gene expression was compared by RNA-Seq **(Figure 1A)**. We found that *S. mutans* exhibited 166 differentially expressed genes (DEGs) when saliva was present **(Supplemental Table 1) (Figure 1B)**. Out of the 166 DEGs, 78 genes were upregulated in response to saliva supplementation **(Figure 1C)**. Notably, we observed increased expression of 12 phosphotransferase system (PTS) genes important for carbohydrate import and utilization in *S. mutans*. Interestingly, we noted significantly increased expression of PTS genes involved in transport of cellobiose (SMU.1596-1600) (27) as well as the PTS specific for glucose/mannose (*manLMN*, SMU.1877-1879) that is a dominant effector for sugar-specific, CcpA-independent catabolite repression (28–30) consistent with our previous study performed in TY (24), supporting a reproducible saliva-associated response across these two different media types. Other notable genes that were upregulated were the global transcriptional regulator *ccpA* (SMU.1591) responsible for carbon catabolite repression in *S. mutans* amongst other roles (31–33), and the *pur* operon (SMU.29-51), responsible for the *de novo* synthesis of purine (34, 35). Interestingly, the *adh* operon (SMU.127-131), which encodes components involved in acetoin catabolism to acetaldehyde and acetyl-CoA (36) was among the most strongly upregulated operons.

**Figure 1.**
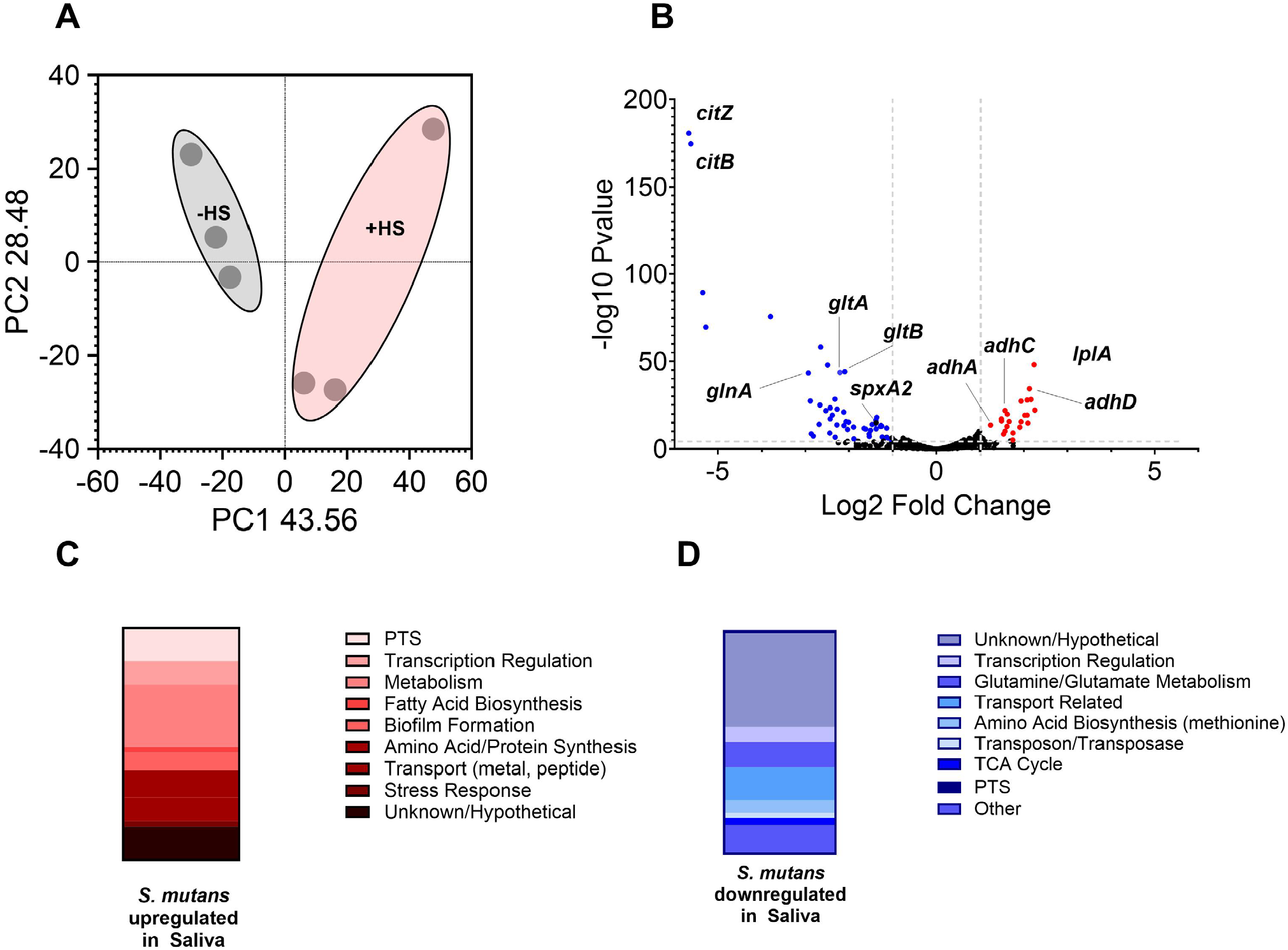
Saliva supplementation alters gene expression in Streptococcus mutans. **(A)** Principal component analysis (PCA) of RNA-Seq profiles from *S. mutans* UA159 grown in CDM without saliva (™HS; gray) or supplemented with 50% (v/v) pooled human saliva (+HS; red). Each point represents an independent biological replicate (n = 3 per condition). PC1 and PC2 explain 43.56% and 28.48% of the variance, respectively. **(B)** Volcano plot showing gene expression changes in +HS relative to ™HS cultures. Each point represents a gene, plotted by log2 fold change and ™log10 (P value). Differentially expressed genes meeting thresholds of log2 fold change ≥ (-)1 and >4 -Log10 P value are shown in red (upregulated) or blue (downregulated); remaining genes are shown in black. Dashed lines indicate the thresholds, and selected genes are labeled. **(C, D)** Stacked bar charts showing the functional classifications of upregulated (C) and downregulated (D) genes. Colors indicate predicted functional categories, as listed beside each chart. PTS, phosphotransferase system; TCA, tricarboxylic acid.

We also documented 88 DEGs that were significantly downregulated in *S. mutans* when grown in saliva **(Figure 1D)**. Thirty-eight of these downregulated genes were annotated as hypothetical genes. Interestingly, among the most strongly downregulated genes were *citB–citZ–idh* (SMU.670-SMU.672) as well as genes involved in glutamine (SMU.363, SMU.364) and glutamate metabolism (SMU.365, SMU.366) (37). We also observed the downregulation of *spxA2* (SMU.2084c), an important transcriptional regulator that coordinates cell envelope homeostasis and oxidative stress response in *S. mutans* (22, 38).

We noted that selected saliva-associated transcriptional changes overlapped with responses previously reported under altered nutrient and stress conditions. Induction of *adh* has been observed during acute glucose-induced acidification, oxygen exposure, and peroxide stress (22, 39, 40), whereas repression of the *citB*– *citZ*–*idh* locus has been reported during acid adaptation and aerobic growth (40–42). These parallels suggest that growth with saliva engages metabolic pathways also responsive to environmental stress; however, the differences among individual genes indicate that the saliva response is distinct from any single previously characterized condition.

### Streptococcus mutans modifies pyruvate-associated metabolite profiles in response to saliva

The transcriptional changes in carbohydrate transport and carbon metabolism prompted us to examine whether saliva addition also altered the metabolite profile of *S. mutans*. Cultures were grown in CDM -HS/+HS to late exponential growth phase (OD_600 nm_ ∼ 1.0), allowing extracellular metabolites to accumulate. Selected metabolites were quantified using colorimetric assays, with extracellular metabolites measured in culture supernatants and acetyl-CoA measured in whole-cell lysates from harvested cells.

We first examined pyruvate and acetyl-CoA, central intermediates connecting carbohydrate metabolism to fermentation and biosynthesis. Pyruvate supplies carbon to the downstream fermentation pathways, whereas acetyl-CoA contributes to acetate and ethanol formation and biosynthetic processes such as fatty acid synthesis. Saliva-supplemented cultures had significantly lower extracellular pyruvate concentrations **(Figure 2A)** and lower cell-associated acetyl-CoA levels **(Figure 2B)** than cultures grown in CDM alone.

**Figure 2.**
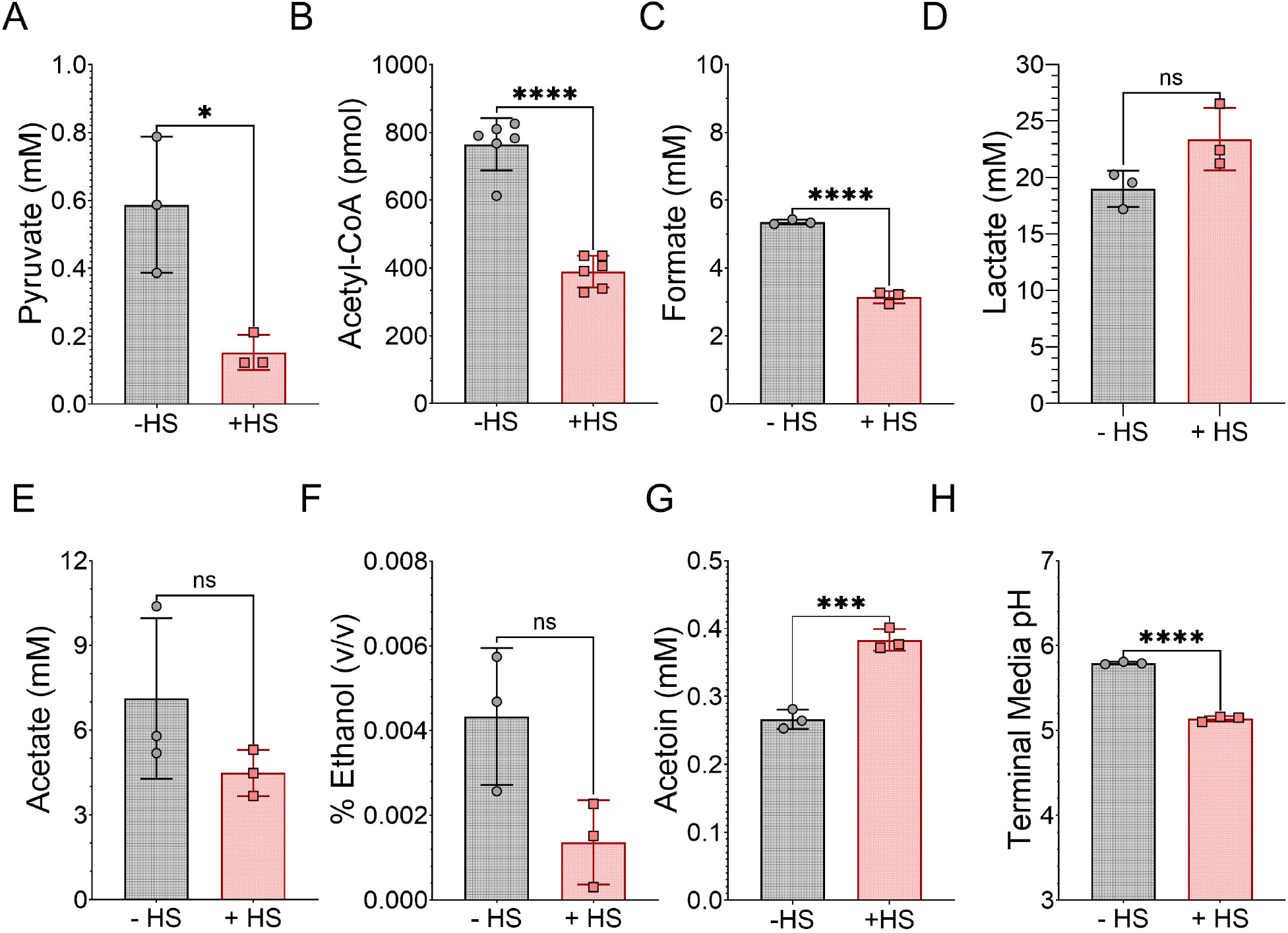
Saliva supplementation alters metabolite profiles and terminal culture pH in Streptococcus mutans. *S. mutans* UA159 was grown in CDM without saliva (™HS; gray circles) or supplemented with 50% (v/v) pooled human saliva (+HS; red squares) to an optical density at 600 nm of approximately 1.0. Quantification of **(A)** pyruvate, **(B)** acetyl-CoA, **(C)** formate, **(D)** lactate, **(E)** acetate, **(F)** ethanol, and **(G)** acetoin. Metabolites were measured in culture supernatants, except acetyl-CoA, which was measured in whole-cell lysates. **(H)** Terminal culture pH. Symbols represent individual measurements of independent biological replicates, n = 3. Differences between ™HS and +HS conditions were assessed using unpaired t-tests. *, P ≤ 0.05; ***, P ≤ 0.001; ****, P ≤ 0.0001; ns, not significant (P > 0.05).

We next quantified lactate, the principal product of homolactic fermentation, and the mixed-acid fermentation products formate, acetate, and ethanol. Formate concentrations were significantly lower in +HS cultures **(Figure 2C)**. Lactate concentrations trended higher in cultures grown with saliva **(Figure 2D)**, while acetate **(Figure 2E)** and ethanol **(Figure 2F)** trended lower, although none of these differences were statistically significant (*P* = 0.08, 0.12, and 0.05, respectively). We also measured the accumulation of extracellular acetoin as the *adh* operon was identified in our transcriptomic dataset and impacts acetyl-CoA and ethanol. Acetoin concentrations were significantly higher in cultures grown with saliva **(Figure 2G)**. This difference, together with the increased expression of the *adh* operon, identified acetoin-associated metabolism as a prominent feature of the *S. mutans* response to saliva.

Lastly, we assessed the terminal pH of cultures as concentrations of several metabolite end products were impacted. We observed significantly lower pH (5.13 ± 0.01) in +HS cultures compared to those grown in CDM alone (5.79 ± 0.01) **(Figure 2H)**, suggesting that *S. mutans* grown in saliva acidifies the environment to a greater extent and may require employment of stress tolerance mechanisms to respond to this environmental pressure.

### Saliva increases acetoin accumulation across Streptococcus mutans strains

The transcriptional and metabolite analyses identified acetoin metabolism as a pathway responsive to saliva addition. Acetoin is synthesized from the condensation of two pyruvate molecules to acetolactate that is subsequently decarboxylated to form acetoin **(Figure 3A)**. The enzymes catalyzing these reactions, acetolactate synthase and acetolactate decarboxylase, are predicted to be encoded by *alsS* and *aldB* (SMU.1452 and SMU.1451), respectively (36). Acetoin can subsequently be catabolized to acetaldehyde and acetyl-CoA by the acetoin dehydrogenase complex encoded by *adhABCD* (SMU.127–130). However, it has not been definitively shown that these genes are involved in acetoin metabolism in *S. mutans*. To investigate the contributions of these pathways to acetoin accumulation during growth with saliva, we constructed a deletion of *alsS*–*aldB* (Δ*alsS*Δ*aldB*) and a nonpolar deletion of *adhC*, which encodes the lipoylated E2 component of the acetoin dehydrogenase complex. We also constructed a nonpolar deletion of *lplA* (SMU.131), encoding a predicted lipoyl ligase implicated in modification of the complex (36, 43, 44). The Δ*lplA* mutant is predicted to retain the structural components of the acetoin dehydrogenase complex but exhibit impaired activity.

**Figure 3.**
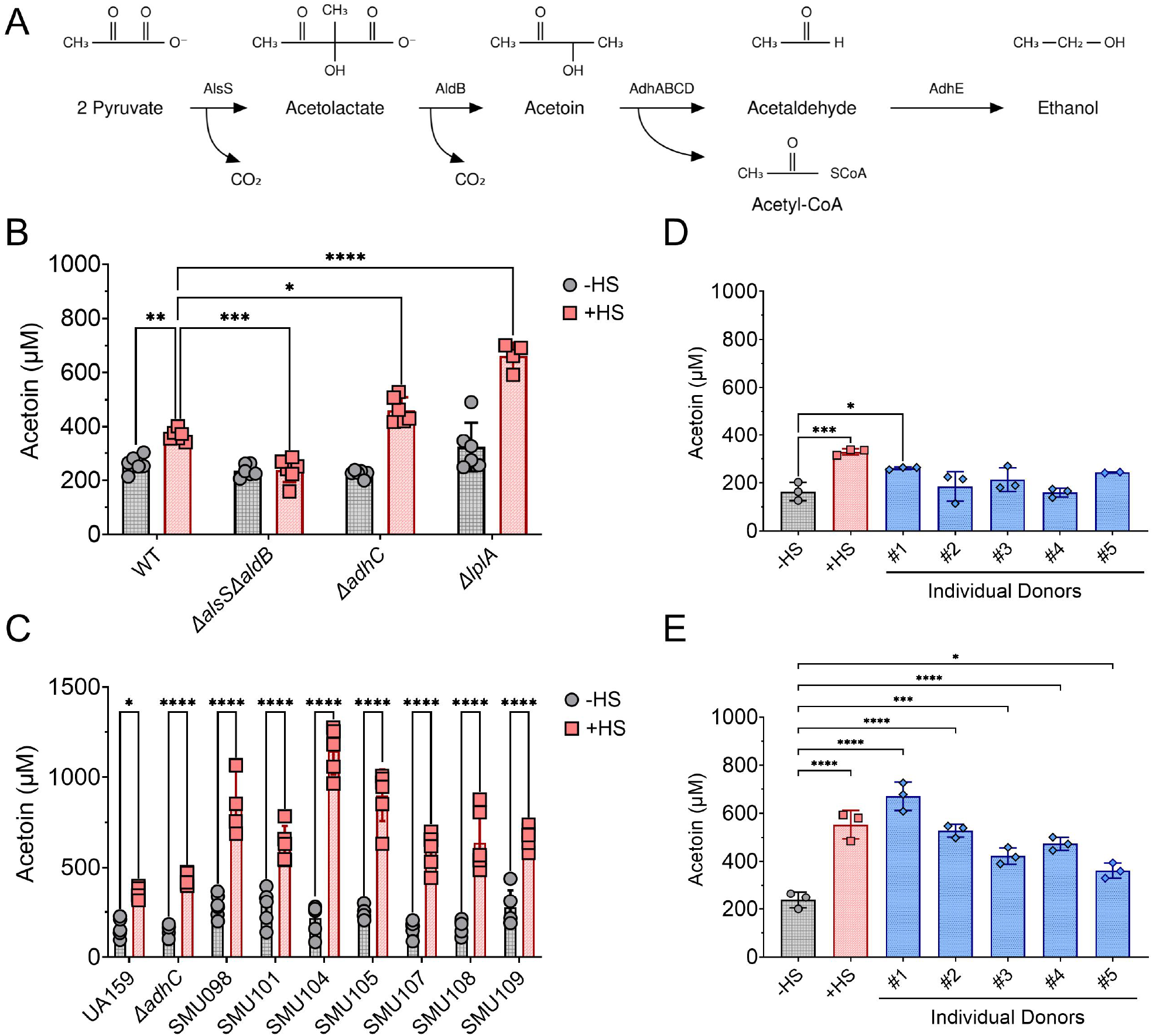
Saliva promotes acetoin accumulation across Streptococcus mutans strains and saliva sources. **(A)** Schematic of acetoin synthesis and catabolism in *S. mutans*, showing the associated enzymes and metabolic intermediates. **(B)** Extracellular acetoin concentrations in cultures of UA159 (WT), Δ*alsS*Δ*aldB*, Δ*adhC*, and Δ*lplA* grown in CDM without saliva (−HS; gray circles) or supplemented with 50% (v/v) pooled human saliva (+HS; red squares). **(C)** Extracellular acetoin concentrations in UA159, the UA159 Δ*adhC* mutant, and the indicated *S. mutans* isolates grown under the same conditions. **(D, E)** Extracellular acetoin concentrations in cultures of UA159 (D) and Δ*adhC* (E) grown in CDM without saliva (gray circles), with pooled human saliva (red squares), or with saliva from five individual donors (blue diamonds; donors #1–#5). All saliva-supplemented media contained 50% (v/v) saliva. Acetoin was quantified in culture supernatants using the Voges–Proskauer reaction. Symbols represent individual measurements of independent biological replicates, n = 3 or 4. Statistical significance was assessed using ordinary two-way ANOVA with Tukey’s multiple comparisons test. Brackets identify the groups compared. *, P ≤ 0.05; **, P ≤ 0.01; ***, P ≤ 0.001; ****, P ≤ 0.0001.

The parent strain UA159 and the deletion mutants were cultured in CDM -HS/+HS, and extracellular acetoin was quantified using the Voges–Proskauer reaction (45). Consistent with our initial metabolite analysis, saliva supplementation significantly increased acetoin accumulation in the isogenic parent strain UA159 **(Figure 3B)**. This increase was abolished in the Δ*alsS*Δ*aldB* mutant, whose acetoin levels remained comparable to those in cultures grown without saliva. Conversely, the Δ*adhC* and the Δ*lplA* mutants accumulated significantly more acetoin than the parent strain with saliva. Taken together, these findings support a requirement for *alsS–aldB* in saliva-associated acetoin accumulation and are consistent with impaired acetoin catabolism following deletion of either *adhC* or *lplA* (36, 46).

To determine whether this response extended beyond the lab-adapted *S. mutans* strain UA159, we quantified acetoin in additional *S. mutans* isolates grown under the same conditions. Saliva supplementation significantly increased acetoin accumulation in every strain tested, with UA159 actually exhibiting the lowest concentration of acetoin among these isolates **(Figure 3C)**. We next examined whether individual donor-sourced saliva reproduced the effects observed with our commercially pooled saliva. Unstimulated saliva collected from five pediatric dental clinic patients (n = 5) was processed using the same procedure as the pooled material. *S. mutans* exhibited similar growth in cultures supplemented with pooled or individual donor-sourced saliva **(Supplemental Figure 1)**, and acetoin accumulation was also observed with individual donor samples **(Figure 3D)**, with a more pronounced increase in the Δ*adhC* mutant **(Figure 3E)**. Thus, the effects of saliva on growth and acetoin accumulation were reproducible using both pooled and individually sourced samples.

Finally, we examined acetoin accumulation in the commensal streptococci *S. gordonii* and *S. sanguinis* grown under the same conditions. Neither species exhibited substantial accumulation with or without saliva **(Supplemental Figure 2)**. Together, these results identify increased extracellular acetoin accumulation as a response shared by the tested *S. mutans* strains and distinguish this response from that of the two other oral streptococcal commensals examined.

### Glucose availability promotes acetoin accumulation during growth with saliva

Because pyruvate serves as the precursor for acetoin synthesis, we investigated whether carbohydrate availability influenced acetoin accumulation during growth with saliva. The wild-type (WT) and the Δ*adhC* mutant were cultured - HS/+HS and supplemented with increasing concentrations of glucose (10, 20, 40, or 80 mM). Extracellular acetoin and terminal culture pH were measured to assess the relationship between glucose availability, acetoin accumulation, and growth medium acidification. Acetoin accumulation generally increased with glucose concentration, with the magnitude of the response varying by strain and whether saliva was present **(Figure 4A)**. In saliva-supplemented cultures, both WT and Δ*adhC* accumulated significantly more acetoin at 40 and 80 mM glucose than at 10 mM glucose, whereas the difference between 10 and 20 mM was not significant. Notably, the Δ*adhC* mutant also exhibited significant increases at 40 and 80 mM glucose in CDM alone. In contrast, the WT strain showed a smaller response without saliva, with neither 40 nor 80 mM glucose yielding significantly greater acetoin accumulation than 10 mM glucose. Across the glucose concentrations tested, acetoin concentrations were higher in +HS cultures than in corresponding -HS cultures, and higher in the Δ*adhC* mutant than in the parental strain, as expected.

**Figure 4.**
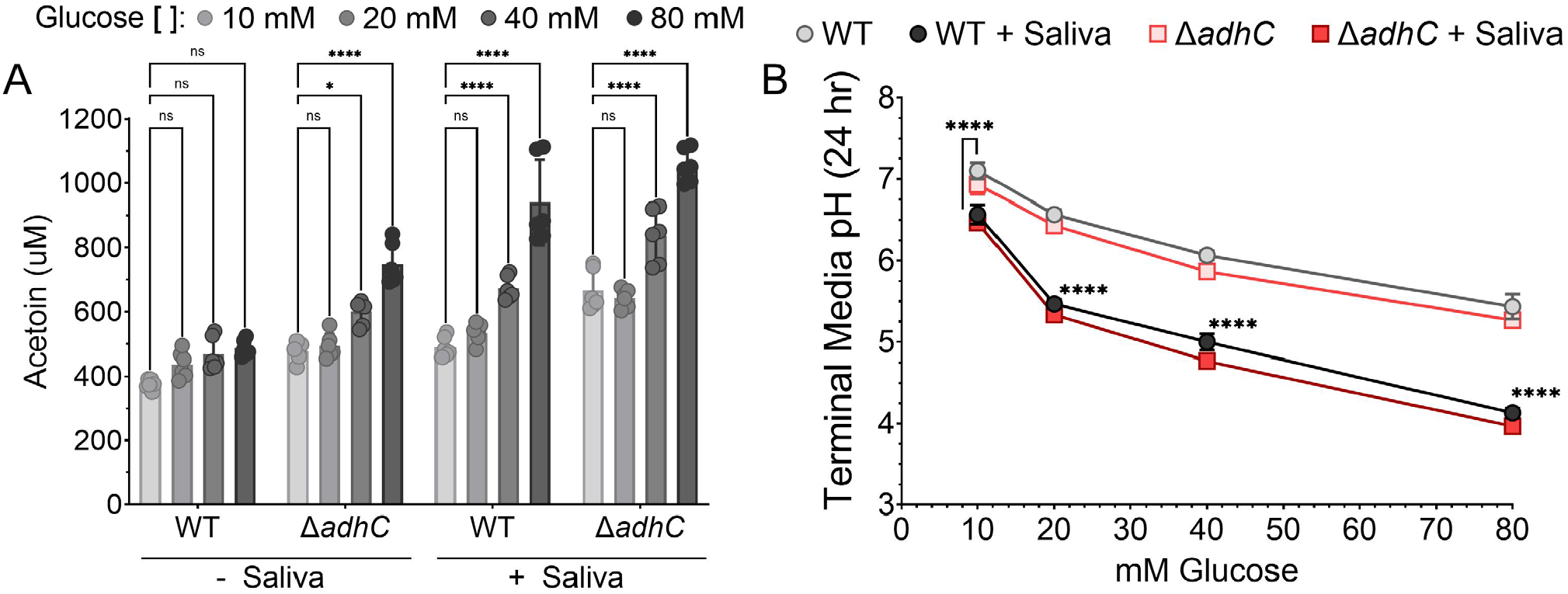
Increasing glucose availability promotes acetoin accumulation and lowers terminal culture pH in Streptococcus mutans. *S. mutans* UA159 (WT) and Δ*adhC* were cultured in CDM containing 10, 20, 40, or 80 mM glucose without saliva (−HS) or supplemented with 50% (v/v) pooled human saliva (+HS). **(A)** Extracellular acetoin concentrations quantified in culture supernatants. Increasing glucose concentrations are represented by progressively darker shades of gray. Symbols represent individual measurements of independent biological replicates, n = 6. Indicated comparisons are between 10 mM glucose and each higher concentration within the corresponding strain and saliva condition. **(B)** Terminal culture pH after 24 hours of growth. WT is represented by circles and Δ*adhC* by squares; lighter gray/red symbols indicate −HS cultures, and darker gray/red symbols indicate +HS cultures. Symbols represent the mean of independent biological replicates, n = 6. Statistical comparisons in panel B were restricted to WT cultures -HS/+HS at each glucose concentration. Statistical significance was assessed using ordinary two-way ANOVA with Tukey’s multiple comparisons test. *, P ≤ 0.05; ****, P ≤ 0.0001; ns, not significant (P > 0.05).

Increasing glucose concentration was accompanied by progressively lower terminal pH in both strains under both media conditions **(Figure 4B)**. Saliva-supplemented cultures reached lower terminal pH values than cultures grown in CDM alone at each glucose concentration. WT and Δ*adhC* exhibited broadly similar acidification patterns, despite their differences in acetoin accumulation. Thus, increased acetoin accumulation and utilization of the acetoin pathway occurred alongside greater medium acidification. Together, these results demonstrate that glucose availability influences acetoin accumulation and that this response is enhanced during growth with saliva. The findings are consistent with acetoin formation providing an alternative outlet for pyruvate derived from carbohydrate metabolism.

### Acetoin metabolism contributes to saliva-associated hydrogen peroxide tolerance in Streptococcus mutans

Previous studies have linked acetoin metabolism to the Spx-regulated oxidative stress response in *S. mutans* (19, 21, 22, 38). Previous work showed that peroxide exposure increases expression of acetoin metabolism genes and acetoin production in *S. mutans* (22). We therefore investigated whether growth with saliva enhanced peroxide tolerance and whether acetoin metabolism contributed to this phenotype. Overnight cultures of WT *S. mutans* and the acetoin pathway mutants were inoculated into CDM -HS/+HS. We then diluted hydrogen peroxide (H_2_O_2_) to a final concentration of 0.03% (8.82 mM) or 0.003% (0.882 mM) and measured their growth over time. At 0.003% H_2_O_2_, growth was comparable to that of untreated cultures across all strains and media conditions (data not shown). At 0.03% H_2_O_2_, however, no measurable growth was detected for any strain in CDM alone. Interestingly, saliva addition supported growth of both WT and the acetoin pathway mutants at this higher peroxide concentration **(Figure 5A–5D)**, demonstrating that saliva provides protection against H_2_O_2_ challenge and that this effect remained following disruption of acetoin metabolism.

**Figure 5.**
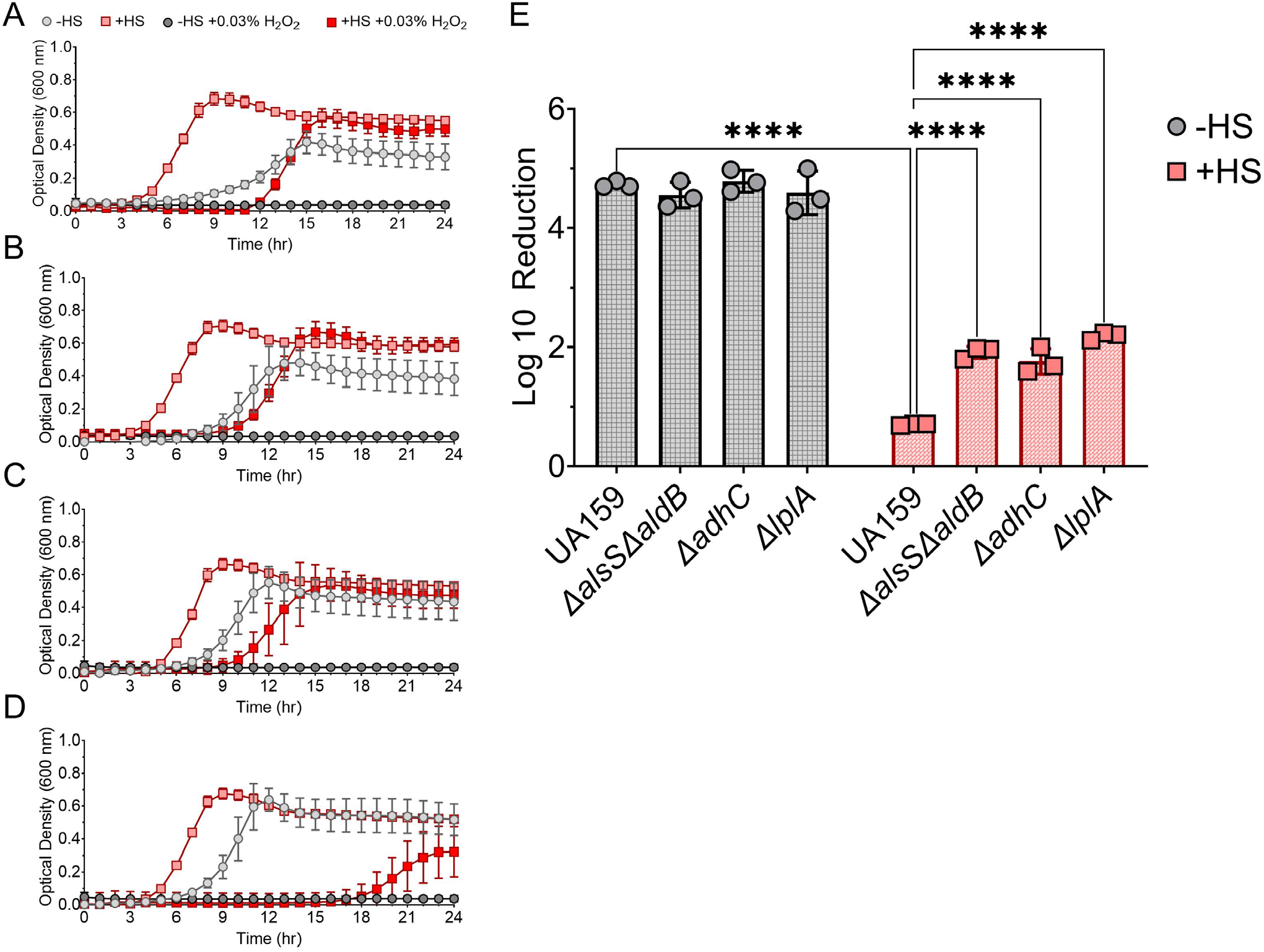
Growth with saliva enhances hydrogen peroxide tolerance in Streptococcus mutans. Growth curves of **(A)** UA159, **(B)** Δ*alsS*Δ*aldB*, **(C)** Δ*lplA*, and **(D)** Δ*adhC* cultured in CDM without saliva (−HS; gray circles) or supplemented with 50% (v/v) pooled human saliva (+HS; red squares), with or without 0.03% H_2_O_2_. Lighter symbols indicate untreated cultures, and darker symbols indicate H_2_O_2_-treated cultures. Growth was monitored by optical density at 600 nm over 24 hours. Symbols represent means of 3 independent cultures. **(E)** Peroxide-mediated killing following growth in CDM without saliva (gray circles) or with saliva (red squares). Cultures were grown to an OD_600_ of approximately 1.0, washed, and challenged in 0.1 M glycine buffer containing 0.5% H_2_O_2_ for 20 minutes. Viability was determined by CFU enumeration and expressed as Log10 reduction; lower values indicate greater survival. Symbols represent individual measurements (n = 3 per group). Statistical significance in panel E was assessed using two-way ANOVA with Tukey’s multiple comparisons test. Brackets identify the groups compared. ****, P ≤ 0.0001.

To quantify peroxide survival independently of growth, WT and mutant strains were grown in cultures of CDM -HS/+HS to an OD_600 nm_ of approximately 1.0. Cells were then washed and challenged in 0.1 M glycine buffer containing 0.5% (147 mM) H_2_O_2_ for 20 minutes. Following growth with saliva, UA159 exhibited approximately 10,000-fold greater survival than cells grown without saliva **(Figure 5E)**. The acetoin pathway mutants (Δ*alsS*Δ*aldB*, Δ*adhC*, and Δ*lplA*) also exhibited increased survival following growth with saliva; however, their survival was approximately 100-fold lower than that of the parent strain. These results indicate that growth with saliva enhances *S. mutans* tolerance to peroxide-mediated killing and that acetoin metabolism contributes to this protection.

### Passaging a pooled Streptococcus mutans transposon library reveals saliva-associated fitness determinants

Our efforts to this point have demonstrated that the acetoin pathway is an important contributor to *S. mutans* physiology when grown in the presence of saliva. However, we sought to develop a more complete picture of other important fitness determinants of *S. mutans* grown with saliva. To this end, we utilized a pooled transposon library of *S. mutans* UA159 that has been used in previous studies (47, 48). The transposon library was initially grown in BHI medium from freezer stocks to an OD_600 nm_ of approximately 1.0. These initial cultures were then inoculated into either CDM (-HS) or CDM supplemented with saliva (+HS). Cultures were transferred into fresh medium every 12 hours for five passages. Samples from the starter culture and passages 1 and 5 under both conditions were retained for Illumina sequencing **(Figure 6A)**, allowing us to assess how the composition of the mutant pool changed between passages across each condition. After mapping reads corresponding to transposon insertion sites back to the *S. mutans* genome, we performed a principal-component analysis (PCA) and observed distinct clustering of our biological replicates (n = 3) by medium condition and passage number **(Figure 6B)**. Moreover, when we compared normalized transposon insertion read counts for the last passage of -HS against +HS, we observed numerous genes whose read counts significantly varied between conditions **(Supplemental Table 2)**. Normalized insert read counts were plotted to visualize differences in genes whose read counts were significantly higher (green) or lower (red) when passaged with media containing saliva **(Figure 6C)**. Notably, we observed several genes identified by Tn-Seq also exhibited differential expression in our RNA-Seq analysis (Figure 1).

**Figure 6.**
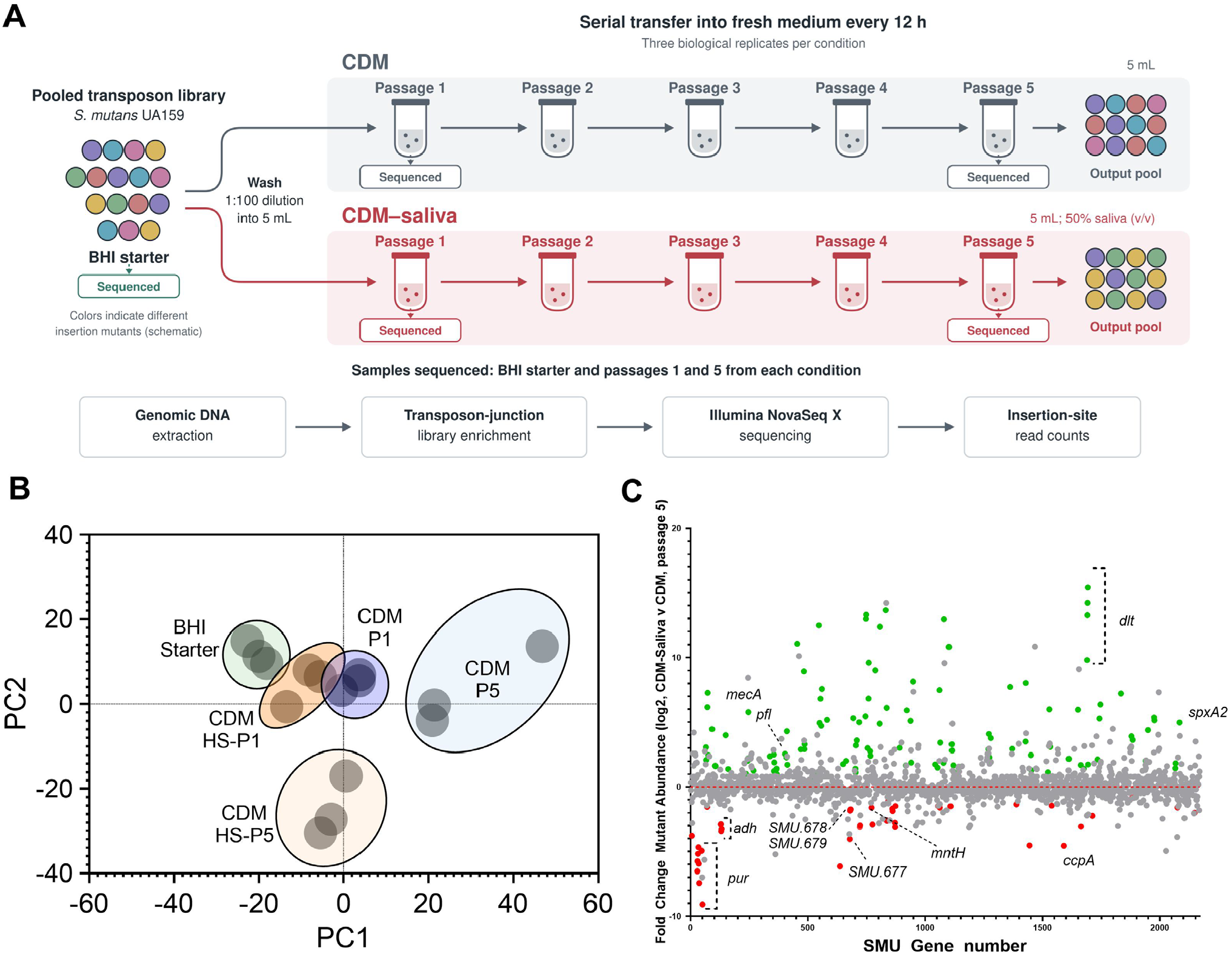
Serial passage of a pooled Streptococcus mutans transposon library identifies saliva-associated fitness determinants. **(A)** Experimental workflow. A pooled *S. mutans* UA159 transposon library grown in BHI was washed and inoculated into CDM without saliva (gray) or supplemented with 50% (v/v) pooled human saliva (CDM HS; red). Cultures were transferred into fresh medium every 12 hours for five passages. Samples from the BHI starter culture and passages 1 and 5 under both conditions were collected for Tn-Seq. Colored cells schematically represent different insertion mutants. **(B)** Principal component analysis (PCA) of gene-level Tn-Seq read counts from the BHI starter culture (green), CDM passage 1 (purple), CDM HS passage 1 (orange), CDM passage 5 (blue), and CDM HS passage 5 (tan). Each point represents an independent biological replicate (n = 3 per group). **(C)** Genome-wide changes in relative mutant abundance following five passages in saliva-supplemented CDM compared with CDM alone. Each point represents the gene-level abundance of mutants carrying transposon insertions, plotted by SMU gene number and log2 fold change. Genes meeting thresholds of log2 fold change ≥ (-)1 and >4 -Log10 P value are shown in green (enriched) or red (depleted); remaining genes are shown in gray. Positive values indicate enrichment with saliva, and negative values indicate depletion. The horizontal dashed line indicates no change, and selected genes and loci are labeled.

Mutants carrying insertions in the *adh* operon were significantly depleted following passage in +HS relative to passage -HS, supporting a contribution of this locus to fitness under saliva-supplemented conditions **(Supplemental Figure 3)**. Further, we saw that both *pur* and *ccpA* were underrepresented in our Tn-Seq dataset and transcriptionally upregulated in our RNA-seq dataset, following a similar trend to the *adh* operon. Interestingly, one of our Tn-Seq hits included *mntH* (SMU.770c) which has been previously identified as a manganese transporter (49), and contrasting with the previously described genes, was found to be transcriptionally downregulated with saliva.

We next wanted to determine how the genes whose insertion mutants were depleted in the Tn-Seq screen contributed to the physiology of *S. mutans* in the presence of saliva, so we constructed mutants of the following identified genes: *purD* (SMU.48), SMU.677, SMU.678, SMU.679, *mntH* (SMU.770c), and *ccpA* (SMU.1591). SMU.677, SMU.678, and SMU.679 are members of a putative three-gene operon. SMU.677 encodes a predicted transcriptional regulator, whereas SMU.678 and SMU.679 are predicted to encode aldo-keto reductase enzymes that reduce carbonyl groups. For these strains, we first analyzed growth in CDM -HS/+HS to determine whether deletion of these genes altered growth patterns or rates. Surprisingly, none of the mutants exhibited significant differences in growth relative to the isogenic parent strain under either condition **(Supplemental Figure 4)**. We also conducted *in vitro* competition experiments between the WT and each mutant strain. The mutant strain was inoculated 1:1 with the isogenic parent in CDM -HS/+HS and grown for 24 hours prior to plating for CFU enumeration. For the majority of the mutant strains tested, we observed no significant change in competitive fitness across either condition examined **(Supplemental Figure 5)**. However, we did see significant changes in fitness in two mutant strains: Δ*mntH* and a double mutant of SMU.678 and SMU.679 (i.e., knockout of both aldo-keto reductase enzymes together). In the Δ*mntH* strain, we observed that addition of saliva increased the competitive fitness of the mutant strain, allowing it to outcompete the WT (∼0.37 log_10_ difference in favor of the mutant) **(Supplemental Figure 5A)**. We observed the opposite phenotype in the double-deletion mutant ΔSMU.678ΔSMU.679, in which the addition of saliva conferred a deleterious effect on the mutant’s competitive fitness (∼0.975 log_10_ difference in favor of the parent) **(Supplemental Figure 5F)**. Interestingly, we observed no changes to competitive fitness in ΔSMU.678 or ΔSMU.677 alone **(Supplemental Figure 5D and 5E)**. These results indicate that depletion of insertion mutants during pooled-library passaging does not necessarily predict a detectable growth defect in monoculture or reduced fitness during pairwise competition with the isogenic parent.

## DISCUSSION

Saliva is a defining component of the oral environment, yet its effects on *S. mutans* metabolism and stress tolerance remain incompletely understood. Here, we show that growth with saliva alters metabolic gene expression and metabolite profiles, with prominent changes in pathways associated with pyruvate metabolism. Increased acetoin accumulation was shared across the *S. mutans* strains examined but was not observed in the two commensal streptococcal species tested. Disruption of acetoin pathway genes also reduced the enhanced peroxide tolerance associated with growth in saliva, linking this metabolic response to stress adaptation. Together, these findings identify acetoin metabolism as an important component of the physiological response of *S. mutans* to saliva and highlight the value of incorporating saliva into models of its behavior in the oral environment.

Acetoin metabolism is often thought to be a means by which fermenting microorganisms can maintain intracellular pH homeostasis, which is important for survival in acidic environments. Indeed, it has been shown that acquisition of acetoin metabolism through genetic manipulation can enhance survival of otherwise sensitive microorganisms at low pH (50). Pyruvate, which is an acidic product formed at the end of glycolysis, has a pKa of ∼2.5, whereas acetoin has a pKa of ∼13.7. This difference in pKa is one of the reasons that acetoin is thought to act as a metabolic “relief valve” for pyruvate, managing carbon overflow while simultaneously mitigating intracellular acid stress caused by glycolytic end products (51). During growth with saliva, increased expression of numerous PTS genes suggested altered carbohydrate uptake capacity in *S. mutans*. Together with the glucose-dependent increase in acetoin accumulation, this observation supports a model in which acetoin formation provides an alternative outlet for pyruvate during carbohydrate-rich growth **(Figure 7A)** (16). During growth with saliva, the Δ*adhC* mutant exhibited significantly higher intracellular pyruvate levels than the parental strain **(Supplemental Figure 6)**. However, the simultaneous accumulation of acetoin and pyruvate in this mutant suggests that acetoin abundance alone does not indicate effective disposal of excess pyruvate. Acetoin turnover, or broader metabolic functions associated with AdhC, may also contribute to maintaining this balance. Although saliva-supplemented cultures reached a lower terminal medium pH, whether growth with saliva alters intracellular pH in *S. mutans* remains to be determined.

**Figure 7.**
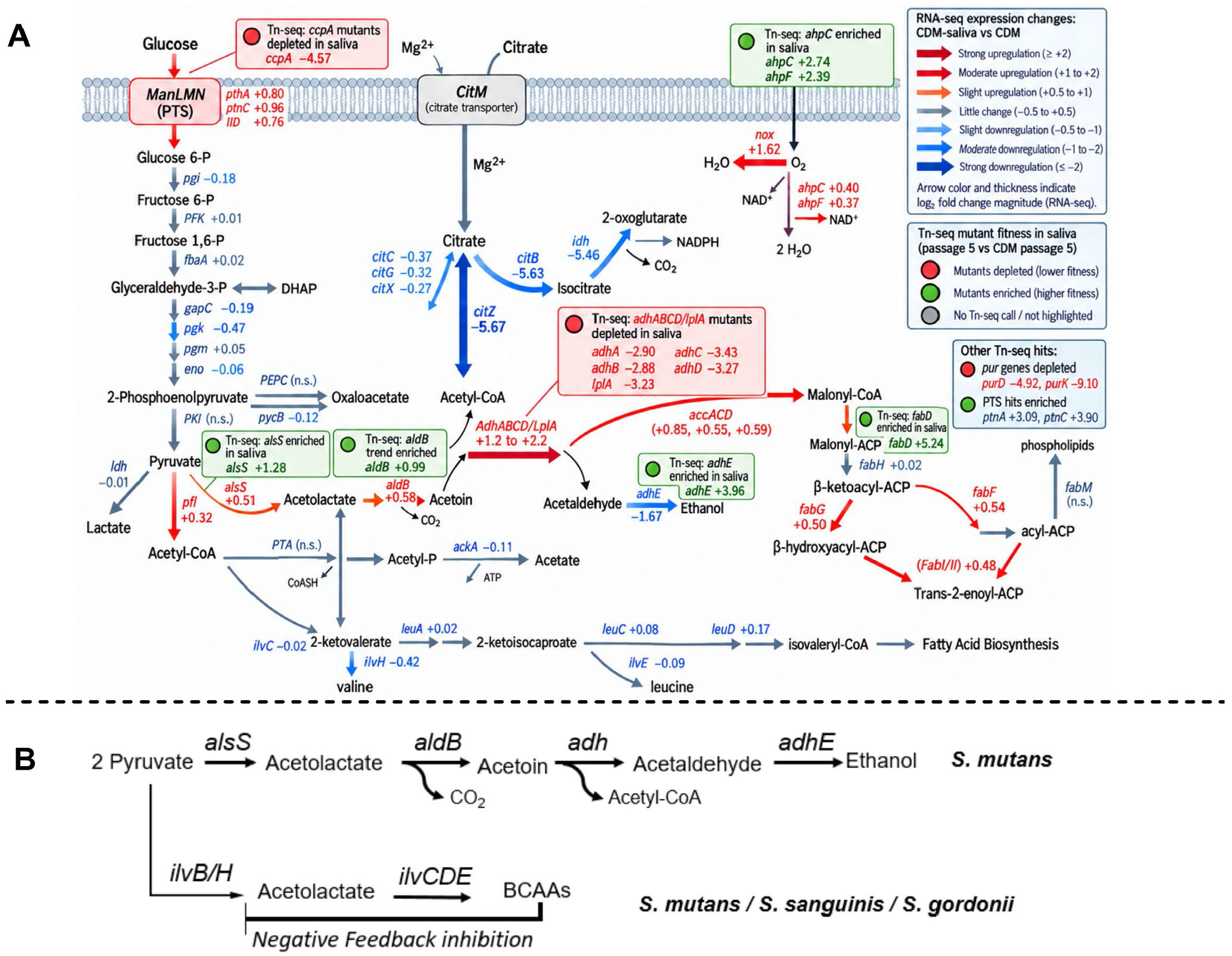
Proposed metabolic responses to saliva and alternative routes of acetolactate formation in oral streptococci. **(A)** Metabolic pathway model integrating RNA-Seq and Tn-Seq results for *S. mutans* grown with saliva. Arrow colors and thicknesses indicate the direction and magnitude of associated gene expression changes in saliva-supplemented CDM relative to CDM alone, as defined in the key. Numbers beside gene names indicate log2 fold changes. Red and green callout boxes identify genes whose insertion mutants were depleted or enriched, respectively, following five passages with saliva relative to CDM alone; values within boxes indicate Tn-Seq log2 fold changes. The nonsignificant enrichment trend for *aldB* is labeled separately. Arrow styling represents transcriptional changes rather than measured metabolic flux. **(B)** Schematic comparison of acetolactate formation through AlsS and IlvBH and its proposed connections to acetoin metabolism and branched-chain amino acid (BCAA) biosynthesis in *S. mutans* UA159, *S. sanguinis* SK36, and *S. gordonii* DL1. The inhibitory line depicts proposed feedback regulation of the biosynthetic route by BCAAs. Pathway connections and regulatory relationships are schematic and have not all been experimentally validated in these species.

Acetoin metabolism has also been suggested to be important for the oxidative stress response in *S. mutans*. Previous studies have shown that the expression of *alsSaldB, adh*, and *lplA* is regulated by SpxA1 (SMU.1142c) and SpxA2 (SMU.2084c), which are critical transcriptional regulators that coordinate responses to oxidative and cell envelope stress in *S. mutans* (19, 21, 22). Mutants carrying transposon insertions in *spxA2* were enriched during growth with saliva, suggesting that disruption of this regulator may confer a fitness advantage under these conditions (Supplemental Table 2). Moreover, previous investigations have shown that SpxA2 may interact directly with AdhC (23), but it remains poorly understood whether this interaction is important to SpxA2 regulation of *adh*. Our findings further support a contribution of acetoin metabolism to oxidative stress tolerance, as acetoin pathway mutants exhibited significantly greater sensitivity to H_2_O_2_-mediated killing following growth with saliva. Acetoin catabolism by the acetoin dehydrogenase complex generates NADH (52), suggesting a potential connection between this pathway and cellular redox balance. We also observed increased expression of *nox* (SMU.1117), which encodes an NADH oxidase that reduces molecular oxygen to water while oxidizing NADH to NAD^+^ (39). These observations raise the possibility that acetoin metabolism contributes to oxidative stress tolerance through effects on NAD^+^/NADH balance. However, we were unable to reliably quantify NADH under the conditions examined using established protocols (53), and this proposed mechanism remains to be tested. Future investigations will seek to understand how acetoin contributes to NAD+/NADH cycling in *S. mutans* and how the presence of saliva may alter it.

Interestingly, when scrutinizing our transposon library data, we found that mutants carrying insertions in the gene responsible for the first step in the formation of acetoin, *alsS*, were enriched in our Tn-Seq experiments. The other gene in the operon, *aldB*, while also enriched, narrowly missed our threshold for being considered statistically significant. This result was initially puzzling when paired with our other observations from the same experiment that demonstrated *adh* was significantly underrepresented, suggesting it is a fitness determinant for *S. mutans* when grown in saliva. We would suspect that *alsS* would also have been similarly underrepresented since we observed increased acetoin accumulation in *S. mutans* cultures grown in the same conditions. However, in another lactic acid bacterium, *Lactococcus lactis*, overproduction of acetoin has been shown to damage DNA and proteins and negatively influence viability *in vitro* (51). Considering this, it is plausible that the *adh* operon is important for *S. mutans* during cultivation with saliva since it is required to effectively catabolize acetoin and failing to do so may be deleterious to *S. mutans*. Nevertheless, our data from our growth experiments would suggest that disruption of *adh* and the resulting increased accumulation of acetoin observed would not be sufficient to impair growth in UA159. Thus, whether acetoin toxicity contributes to the pooled-library fitness phenotype remains unresolved.

The increased activity of the acetoin pathway observed in *S. mutans* raised the question of how other important oral *Streptococcus* species may utilize this pathway and how it compares to *S. mutans*. Under the conditions tested, we found that two important commensal oral streptococci, *S. sanguinis* (SK36) and *S. gordonii* (DL1), exhibited little detectable acetoin accumulation in CDM - HS/+HS and showed acetoin-associated signals comparable to those of our Δ*alsS*Δ*aldB* operon deletion strain that lacks acetoin synthesis ability (Supplemental Figure 2). It should be noted that our acetoin quantification experiments utilized Voges-Proskauer chemistry, which relies on conversion of acetoin to a diacetyl intermediate prior to condensing with the other reactants to form a red-pink product that is measured spectrophotometrically. Consequently, diacetyl that is naturally produced by these species causes modest but detectable background noise in our results, which is our primary hypothesis for why we see a low residual assay signal in our Δ*alsS*Δ*aldB* mutant.

Regardless, we noted that both *S. sanguinis* and *S. gordonii* encode genes that are annotated as acetolactate synthase; however, these genes fall within the *ilv* operon responsible for isoleucine and valine biosynthesis in these species (52). In fact, similar to acetoin, the initial step in the synthesis of L-valine is the condensation of two pyruvate molecules to form acetolactate, a reaction catalyzed by the acetolactate synthase encoded by *ilvBH* (SSA_1969-1970 ; SGO_0526-0527). Additionally, orthologous genes exist (SMU.231-232) within *S. mutans* that catalyze this same process (54), meaning *S. mutans* encodes two enzyme systems predicted to catalyze the same reaction. However, the acetolactate synthase encoded by *ilvBH* is distinct from AlsS because it comprises both a large catalytic subunit (IlvB) and a small regulatory subunit (IlvH) that is subject to multivalent repression by valine and isoleucine that prevents the transcription of *ilvB* via ribosome-mediated transcriptional attenuation (55–57). As a result, acetolactate production from *ilvB* is limited in the presence of these branched-chain amino acids (BCAAs) **(Figure 7B)**. Conversely, *alsS* appears to only encode the catalytic protein and not the regulatory sequence seen in *ilvBH*. Therefore, *S. mutans* is likely able to source acetolactate from two enzymatic routes whereas *S. gordonii and S. sanguinis* can only source acetolactate from *ilvB*. However, this has yet to be experimentally validated in these species and future investigation is required to develop a more comprehensive understanding of these processes across oral streptococci. Nevertheless, it is noteworthy that *S. mutans* appears to have acquired multiple means to produce acetolactate that is not tethered to the presence of valine or isoleucine and this seemingly differs from other oral *Streptococcus* species.

The increased tolerance *S. mutans* exhibited to H_2_O_2_ when cultured in saliva was an intriguing finding of this investigation. In our preliminary efforts to understand this phenotype, we were able to identify that acetoin metabolism was only a partial contributor to the increased H_2_O_2_ tolerance observed, likely due to the multifactorial nature of the oxidative stress response of *S. mutans*. For example, during our assessment of changes in the *S. mutans* transcriptome during growth with saliva, we observed a significant downregulation of the *citB-citZ-idh* locus, which is part of the *S. mutans* incomplete TCA cycle, consistent with previous studies that examined the *S. mutans* transcriptome during oxygen stress (39, 42). The initial step in the incomplete TCA cycle of *S. mutans* is catalyzed by citrate synthase, encoded by *citZ*, which condenses acetyl-CoA and oxaloacetate to form citrate and coenzyme A (58). Our data showed that *S. mutans* has significantly lower cell-associated acetyl-CoA levels when grown in the presence of saliva. Given both the downregulation of *citZ* and the decrease of acetyl-CoA, we hypothesize that acetyl-CoA may be redirected toward another pathway. During growth with saliva, expression of genes encoding acetyl-CoA carboxylase (SMU.1734–SMU.1736), which converts acetyl-CoA to malonyl-CoA, increased. Genes in the *fab* operon involved in subsequent steps of *de novo* fatty acid biosynthesis were also upregulated (59, 60). Together, the reduced expression of the citrate synthase locus and increased expression of fatty acid biosynthesis genes raise the possibility that growth with saliva redirects acetyl-CoA toward fatty acid synthesis. However, metabolite pool sizes and transcript levels alone do not establish increased flux through this pathway. Indeed, previous reports have shown that *S. mutans* modifies its membrane in response to stressors such as lower pH, which we observed in saliva, or in response to oxidative stress (60). It has also been documented that *S. mutans* increases the ratio of unsaturated fatty acids (UFA) to saturated fatty acids (SFA) as a means to mitigate acid stress (61). Deletion of *fabM*, required for UFA production in *S. mutans*, has been shown to increase sensitivity to both hydrogen peroxide and acid stress (60, 61). Therefore, it is plausible that in response to saliva, *S. mutans* modifies its cell membrane to tolerate stressors such as reduced pH or various reactive oxygen species. Future investigations will determine whether growth with saliva alters the ratio of unsaturated to saturated fatty acids in *S. mutans* and whether these changes contribute to stress tolerance.

Overall, this study demonstrates that supplementing laboratory growth media with saliva alters transcriptional and metabolic activity in *S. mutans*, revealing a contribution of acetoin metabolism to saliva-associated stress tolerance. Saliva continually bathes supragingival communities, yet most laboratory studies examine oral bacteria in conventional media that lack this host-derived component. The distinct phenotypes observed during growth with saliva underscore the importance of culture conditions in determining bacterial physiology. These findings advance our understanding of how *S. mutans* responds to host-associated environmental factors and highlight the value of incorporating saliva into laboratory models of oral microbial physiology.

## MATERIALS AND METHODS

### Preparation of Bacterial Cultures and Strains

All bacterial strains were resuscitated from frozen glycerol stocks stored at -80°C onto Brain Heart Infusion (BHI) agar plates for at least 48 hours at 37°C and 5% CO_2_, with mutant strains expressing an erythromycin-resistance gene (*ermB*) streaked onto BHI agar supplemented with erythromycin (0.01 mg mL^-1^). To establish planktonic overnight cultures, single, isolated colonies were aseptically transferred to 2 mL of BHI and were grown ∼18 hr at 37°C and 5% CO_2._ Overnight cultures were harvested via centrifugation, washed, and normalized to an optical density at 600 nm of 0.2 in 1x phosphate-buffered saline (PBS) prior to dilution (1:100) into experimental growth media. Bacterial strains used in this study **(Supplemental Table 3)** were cultivated in a chemically defined medium (CDM) specific for streptococci (25, 26).

### Human Saliva and Growth Media Preparation

Commercially sourced pooled whole human saliva was purchased from Innovative Research. The saliva was received frozen and stored at -20°C prior to processing for use. Frozen saliva was thawed overnight at 4°C and centrifuged at 4500 RPM for 10 minutes before being filter-sterilized via passage through a syringe-driven 0.22 μm polyethersulfone (PES) filter unit. 50 mL aliquots were locally stored at - 20°C prior to experimental use and thawed one day prior to use at 4°C. CDM-Saliva (+HS) was prepared via addition of 50% saliva volume by volume (v/v) with CDM that was prepared without the addition of glucose. After mixing CDM and saliva, glucose was then diluted from a 1 M stock into both -HS and +HS, respectively, to a final concentration of 20 mM. Following addition of the glucose, each medium type was filter-sterilized again through a PES syringe-driven filter unit prior to use for bacterial cultivation.

Saliva samples from individual donors were obtained at the Nationwide Children’s Hospital (NCH) Dental Clinic in Columbus, OH, under an institutional review board–approved protocol (IRB #STUDY00003999). Participants were 6– 17 years of age, had an American Society of Anesthesiologists (ASA) physical status classification of I or II and had not used antibiotics or antifungals within the preceding 30 days. Both caries-free and caries-active individuals were included. Unstimulated saliva samples (∼2 mL) were collected by a pediatric dentist, refrigerated until processing and processed in a manner similar to that used for the pooled saliva sample. Written informed consent was obtained from each participant’s guardian and assent was obtained from participants older than 9 years. All procedures were conducted in accordance with relevant institutional guidelines and regulations.

### Construction of Bacterial Mutant Strains

Bacterial mutant strains (Supplemental Table 3) were created via PCR ligation mutagenesis following a previously published method (68). Briefly, 0.1 μg of PCR ligation product containing an erythromycin resistance gene (*ermB*) (62, 69) was transformed into *S. mutans* strain UA159 by induction of competence through treatment with synthetic competence stimulating peptide (sCSP) with BHI medium. After approximately 3 hours of incubation at 37°C and 5% CO_2_, *S. mutans* was plated onto BHI agar plates supplemented with 0.01 mg mL^-1^ erythromycin. The resulting mutants were confirmed to contain no unintended mutations (i.e., SNPs or other genomic rearrangements) through whole genome sequencing (SeqCenter). The primers used for construction of all mutant strains are listed in **Supplemental Table 4**.

### Bacterial Growth Measurements

Overnight cultures were harvested by centrifugation, washed, and normalized to OD_600 nm_ = 0.2 with PBS prior to inoculation (1:100) into the experimental medium of choice. Growth measurements were completed using a Bioscreen C MBR automated turbidometric analyzer (Growth Curves Ab Ltd., Helsinki, Finland) with the OD_600 nm_ recorded every half hour for 24 hours. Wells were overlaid with 0.05 mL sterile mineral oil.

### RNA Isolation and Sequencing

Select bacterial strains were grown in respective growth media until a measured optical density (OD_600 nm_) of ∼0.6. Bacterial cells were harvested by centrifugation and cell pellets were resuspended into 1 mL of RNAprotect (Qiagen). RNA isolation was performed following previously established methods (62). RNA was sequenced through SeqCenter using an 8-million single-read package applied to RNA. Delivered FASTQ files were analyzed by a custom workflow that included FASTQ Groomer (v 1.1.5), FASTQ Quality Trimmer (v 1.1.5), mapping of reads with Bowtie2 (v.2.3.4.3), and htseq-count (v 0.9.1) on genome features from an *S. mutans* UA159-specific GFF3 file (accession NC_004350.2) that resulted in a CSV file containing non-normalized read counts. All read counts from all samples were combined into a single CSV file and uploaded to Degust (63) and edgeR analysis was performed to determine log2 fold change and false discovery rate (FDR) for +HS compared to the -HS condition. Total read counts and alignment data from RNA-seq libraries can be found in **Supplemental Table 5**.

### Pooled Transposon Library Preparation and Sequencing

The *S. mutans* UA159 pooled transposon library used in this study was provided courtesy of Dr. Robert Shields and construction of the library is detailed elsewhere (47, 48). The pooled library was revived from frozen -80°C glycerol stocks through aseptic transfer of approximately 10 μL of frozen stock to 5 mL of BHI broth supplemented with spectinomycin and grown overnight at 37°C and 5% CO_2._ The overnight cultures were washed and normalized to OD_600 nm_ = 0.2 in 1x PBS. The washed culture was inoculated at a dilution of 1:100 into 5 mL of CDM -HS/+HS. The cultures were incubated for 12 hours at 37°C and 5% CO_2_ between passages. For passaging, 1 mL of culture was removed, washed, and used to inoculate 5 mL of fresh medium. The remaining 4 mL of culture was saved for genomic extraction and sequencing. Genomic extraction and preparation of the library were performed following methods detailed previously (47, 64).

Briefly, genomic DNA was isolated via a commercial kit (Qiagen, DNeasy PowerLyzer Microbial Kit) prior to being digested with MmeI for 4 hours at 37ºC followed by heat-inactivation at 65 ºC for 15 minutes. The restriction digest products were purified via phenol:chloroform:isoamyl alcohol extraction (25:24:1) followed by precipitation with isopropanol. The precipitant was pelleted via centrifugation at 10,000 x g for 10 minutes at 4 ºC and washed twice with ice-cold 70% ethanol. The residual ethanol was evaporated via air-drying and the DNA pellet was resuspended in 0.1 mL of nuclease-free water. Barcoded adapter ligation was conducted overnight at 16 ºC using a T4 DNA Ligase (400 U μL^-1^, New England Biolabs). The ligation products were PCR-amplified prior to separation on a 2% TAE agarose gel stained with SYBR Safe Gel Stain. Bands that were approximately 120 bp were excised and purified via QIAquick Gel Extraction Kit following the manufacturer’s protocol before elution with 50 μL nuclease-free water. The DNA yield was quantified via Qubit dsDNA-HS Assay kit. The purified PCR product was validated via TapeStation (High Sensitivity D1000) to confirm band size and concentration. Following this, samples were pooled and normalized to 1 ng μL^-1^ prior to sequencing.

The pooled samples were sequenced through SeqCenter using 1 lane of an Illumina NovaSeq X Plus (2.5 billion, 151 bp paired-end reads). The PhiX internal Illumina Sequencing control was used to enhance base-calling accuracy that can be negatively influenced by the low sequence diversity typical of pooled transposon library samples.

### Tn-Seq Analysis

Tn-Seq sequencing data were processed using the Galaxy platform (65). Sequencing files were concatenated as needed and demultiplexed using sample-specific 5′ barcodes, allowing no barcode mismatches or deletions. Read quality was assessed using FastQC v0.12.1 following processing with FASTQ Groomer. Reads were trimmed using the Galaxy Trim sequences tool and converted to FASTA format. The Mag6 sequence (5′-ACAGGTTGG-3′) was subsequently removed using Cutadapt v4.9, configured for 3′ adapter trimming with a maximum error rate of 0.1 and a minimum overlap of three nucleotides. Reads lacking the adapter or shorter than one nucleotide after trimming were discarded. Retained reads were aligned as single-end reads to the *Streptococcus mutans* UA159 reference genome (accession NC_004350.2) using Bowtie2 (66). Aligned reads were assigned to annotated genes using htseq-count (67) and the corresponding genome annotation file, and gene-level counts were compiled across three biological replicates of the starter cultures and passages 1 and 5 under each growth condition (-HS/+HS). Total read counts and alignment data from each Tn-Seq library can be found in **Supplemental Table 6**.

### Metabolite and pH Quantification

*S. mutans* cultures were grown to an OD_600 nm_ = 1.0 prior to centrifugation at 4500 rpm for 10 minutes at 4ºC to pellet cells. The supernatant was extracted and used for quantification of all metabolites listed excluding acetyl-CoA and intracellular pyruvate, which utilized whole-cell lysates prepared from cell pellets. Quantification of lactate (MAK329), acetate (MAK474), formate (MAK491), ethanol (MAK481), pyruvate (MAK332), and acetyl-CoA (MAK566) was performed using commercially sourced colorimetric kits following the manufacturer’s protocol (Millipore Sigma). Acetoin was quantified via a modified Voges-Proskauer method that has been described previously (45). Briefly, 0.2 mL of sample or known acetoin standard (Sigma, 40127-U) was dispensed into 96-well deep-well polypropylene plates followed by 0.14 mL of 0.5% creatine, 0.2 mL of 5% 1-napthol, and 0.2 mL of 40% potassium hydroxide added sequentially. The reaction was incubated for 15 minutes at room temperature before measuring absorbance (OD_570 nm_). Acetoin was quantified from unknown samples by plotting absorbance values on a standard curve generated from known standards. Terminal medium pH was measured via an electrode from collected supernatants.

### Competitive Index Assays Using Colony Forming Units (CFUs)

1 × 10^5^ cells mL^-1^ each of UA159 (pPGS; spectinomycin resistant) and mutant strains (*ermB*; erythromycin resistant) were inoculated in equal volumes (1:100) into 1 mL of test media. Portions of each inoculum were serially diluted and plated onto both BHI agar plates containing spectinomycin (selection of WT *S. mutans* UA159) and plates containing erythromycin (selection of mutants). Colony forming units (CFUs) were later enumerated from these agar plates after incubation to determine the initial cell count at the time of inoculation (*t*_i_ = 0 h). The remaining inoculum was incubated at 37°C and 5% CO_2_ for 24 hours. Resulting cultures were harvested, washed and resuspended with 1x PBS while transferring to a 5 mL polystyrene round-bottom tube. To isolate single cells, tubes were sonicated within a water bath sonicator (four intervals of 30 seconds, resting 2 minutes on ice). Cultures were serially diluted and plated on both BHI spectinomycin and erythromycin agar plates and incubated for 48 h at 37°C and 5% CO_2_. Following CFU counting of the final cell count (t_f_ = 24 h), a competitive index was calculated using the following formula: CI= Log10[(t_f_ UA159 CFU/t_f_ mutant CFU)/(t_i_ UA159 CFU/t_i_ mutant CFU)].

### Hydrogen Peroxide Challenge Experiments

Hydrogen peroxide stock (30%) was diluted to 0.03% or (1:1000) or 0.003% (1:10,000) into inoculated test media prior to being subjected to growth measurement as described above. The CFU assay was conducted using a previously detailed protocol (70). Briefly, 5 mL bacterial cultures were grown to an optical density of 1.0 (OD_600 nm_) before 1 mL was removed and the cells were harvested via centrifugation (14,000 x g, 1 minute). The supernatant was removed and the cell pellet was resuspended in 1 mL of 0.1 M Glycine buffer. The cell pellets were washed two times before 10 μL was removed for initial CFU enumeration (CFU_i_). The pellet was then resuspended in 1 mL of 0.1 M glycine buffer supplemented with 0.5% hydrogen peroxide. The suspension was incubated for 20 minutes at 37ºC and 5% CO_2_ before 10 μL of sample was removed for final CFU enumeration (CFU_f_). The samples were serially diluted in PBS and plated onto BHI agar and incubated at 37ºC and 5% CO_2_ for 48 hours prior to enumeration of CFUs and calculation of CFU/mL.

### Graphing and Statistics

Graphing of data was completed with GraphPad Prism (version 10.6.1). All statistical analysis were completed with GraphPad built-in analysis tools including PCA of RNA-seq and Tn-Seq data, one-way/two-way ANOVA, and unpaired t-tests.

## DATA AVAILABILITY

The resulting RNA-Seq raw sequencing and data files from this study are available from NCBI-GEO (Gene Expression Omnibus) under accession number **GSE346581**. Sequencing reads for transposon library experiments have been deposited in the Sequence Read Archive (SRA) under the accession number **PRJNA1525691**. The raw and processed data generated in this study have been deposited in FigShare and are publicly available under DOI **10.6084/m9.figshare.34701867**.

## ACKNOLWEDGEMENTS

Research reported in this publication was supported by the National Institute of Dental & Craniofacial Research of the National Institutes of Health under Award Number **5T32DE032994**. The content is solely the responsibility of the authors and does not necessarily represent the official views of the National Institutes of Health.

## AUTHOR CONTRIBUTIONS

Jacob Harris: Investigation, Formal analysis, Writing – original draft.

Alyssa N. Deever: Formal analysis, Data curation, Writing – review & editing. Kyulim Lee: Formal analysis, Data curation, Supervision, Writing – review & editing.

Justin R. Kaspar: Funding acquisition, Project administration, Supervision, Writing – review & editing.

## ARTIFICIAL INTELLIGENCE (AI) UTILIZATION STATEMENT

The authors acknowledge the use of ChatGPT (GPT-6.1 Sol) in the final editing phases to assist with grammar polishing and improving the readability of the manuscript. In addition, ChatGPT (GPT-5.6 Sol) was used to help redesign figures 3A, 6A and 7A in the manuscript that were based on the authors’ own initial versions that were constructed in Microsoft PowerPoint. The authors reviewed and edited the output and take full responsibility for the final content.

## DECLARATION OF CONFLICTING INTERESTS

The authors declare no potential conflicts of interest with respect to the research, authorship, and/or publication of this article.

## Notes

https://doi.org/10.6084/m9.figshare.34701867

https://www.ncbi.nlm.nih.gov/geo/query/acc.cgi?acc=GSE346581

https://www.ncbi.nlm.nih.gov/bioproject/?term=PRJNA1525691

### Competing Interest Statement

The authors have declared no competing interest.

