## Supplemental Figure 1 for "Human Saliva Induces Acetoin-Dependent Metabolic Adaptation that Promotes Stress Tolerance in *Streptococcus mutans*"

**Table of Contents:**

Supplemental Figures (S1-S6): Pages 2 - 9

Supplemental Tables (Table S1-S6): Pages 10 – 26

Supplemental References: Page 27

Supplemental Excel Files: Contains whole genome RNA-Seq and Tn-Seq data (log2 fold change and –log10 P values) used to construct Supplemental Tables 1 and 2.

### Corresponding author

**Mailing address:**
Division of Biosciences, The Ohio State University, College of Dentistry,
305 W. 12th Avenue, Postle Hall Rm 4185, Columbus, OH 43210.


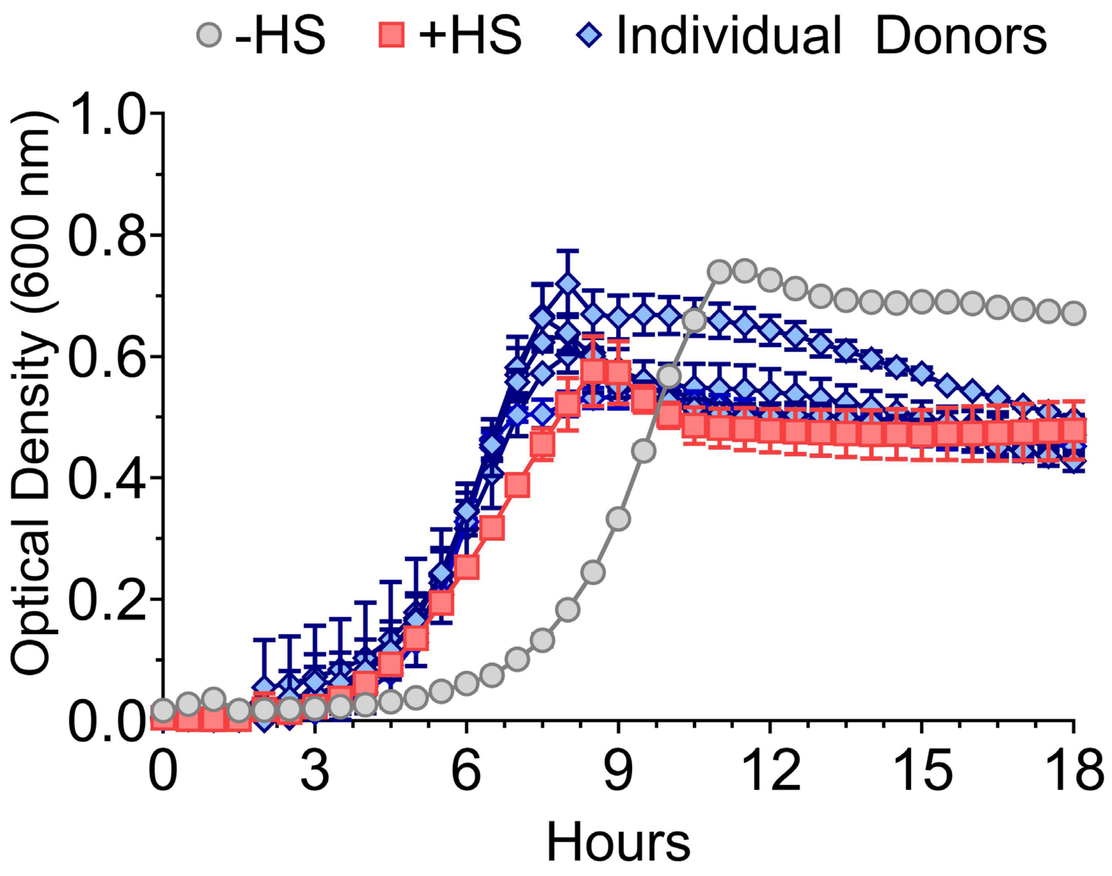

**Supplemental Figure 1. *Effects of pooled and individual-donor saliva on Streptococcus mutans growth.*** Growth curves of *S. mutans* UA159 cultured in CDM without saliva (−HS; gray circles), with pooled human saliva (+HS; red squares), or with saliva from five individual donors (blue diamonds). Saliva-supplemented media contained 50% (v/v) saliva. Growth was monitored by optical density at 600 nm over 18 hr. Each blue curve represents a separate donor (n=5).

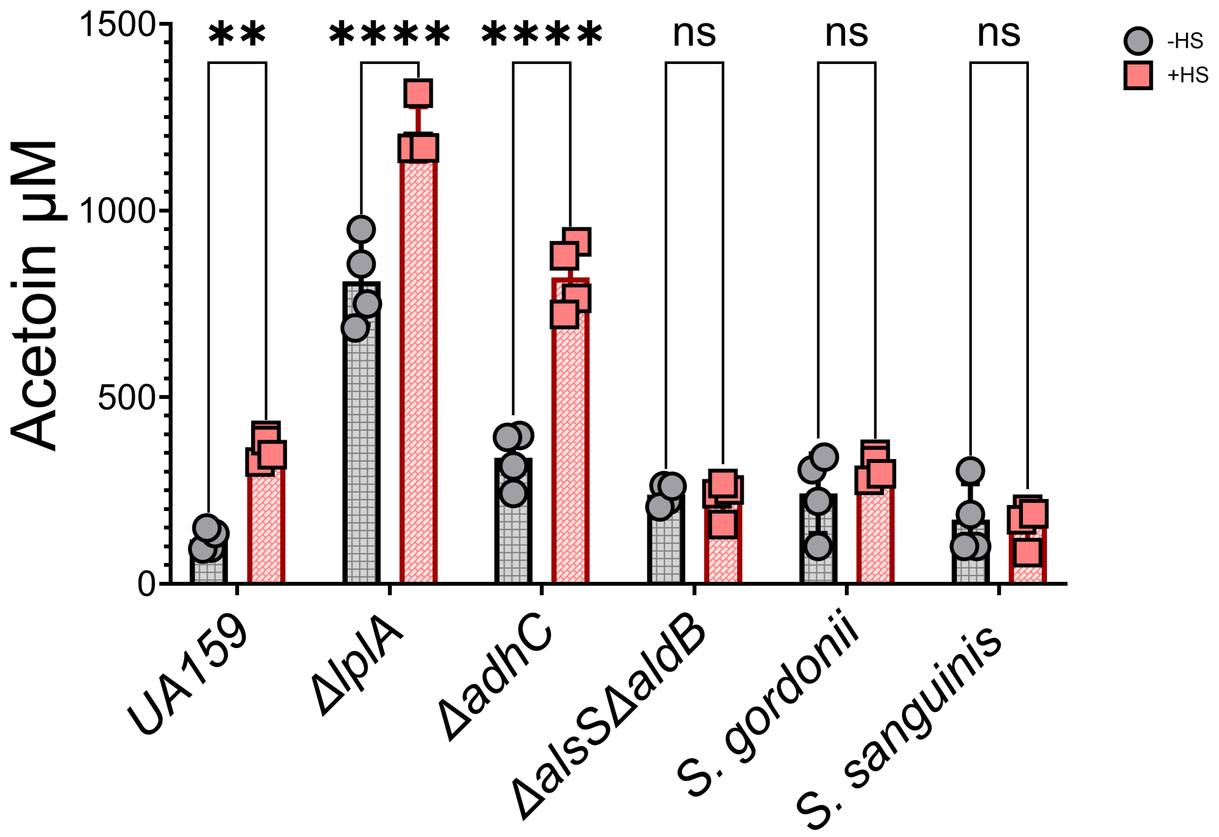

**Supplemental Figure 2. *Acetoin accumulation in Streptococcus mutans and commensal oral streptococci*.** Extracellular acetoin concentrations in cultures of *S. mutans* UA159 and respective Δ*lplA*, ΔadhC, and Δ*alsS*Δ*aldB* mutants, *Streptococcus gordonii* DL1, and *Streptococcus sanguinis* SK36 grown in CDM without saliva (−HS; gray circles) or supplemented with 50% (v/v) pooled human saliva (+HS; red squares). Acetoin was quantified using the Voges–Proskauer reaction. Symbols represent individual independent cultures (n = 4). Statistical significance between −HS and +HS cultures within each strain was assessed using two-way ANOVA followed by Tukey's multiple-comparisons test. **, P ≤ 0.01; ****, P ≤ 0.0001; ns, not significant (P > 0.05).

A

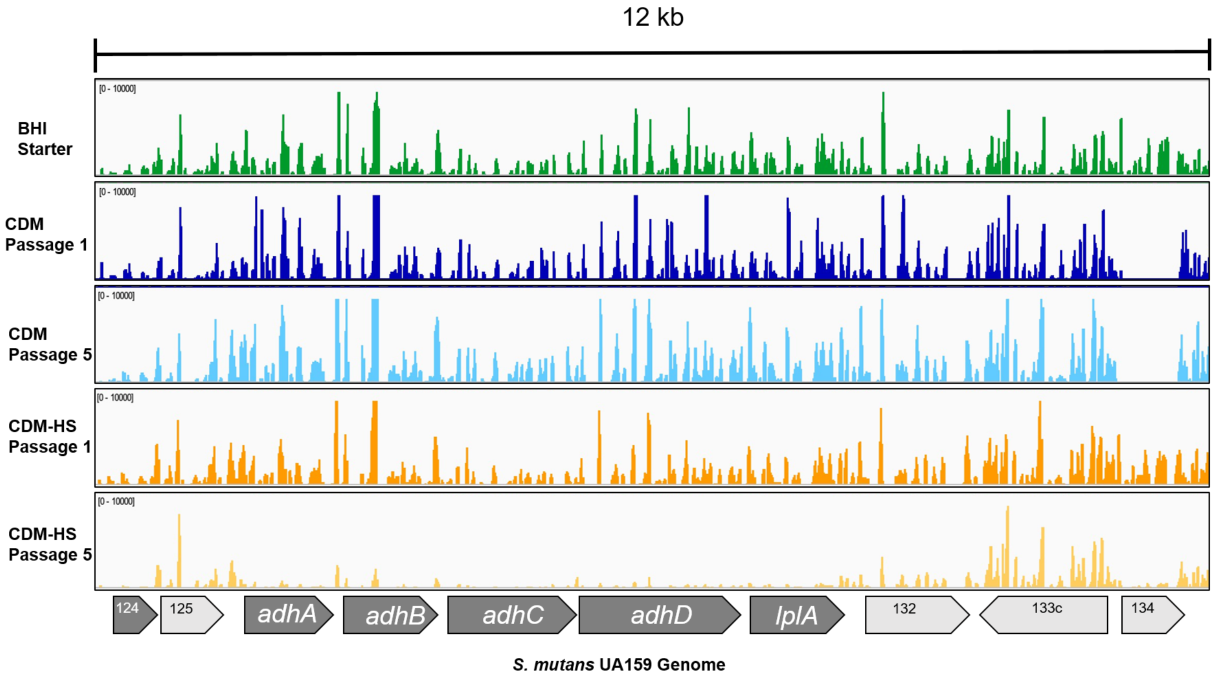
12 kb
BHI Starter
CDM Passage 1
CDM Passage 5
CDM-HS Passage 1
CDM-HS Passage 5
124
125
adhA
adhB
adhC
adhD
lplA
132
133c
134
*S. mutans* UA159 Genome

B

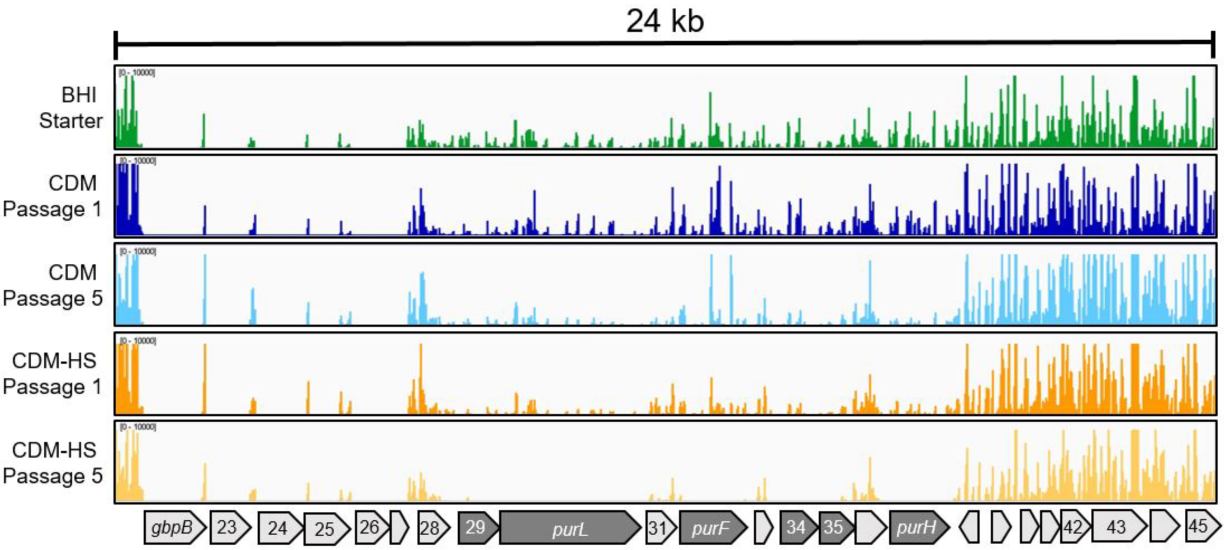
24 kb
BHI Starter
CDM Passage 1
CDM Passage 5
CDM-HS Passage 1
CDM-HS Passage 5
gbpB
23
24
25
26
28
29
purL
31
purF
34
35
purH
42
43
45

C

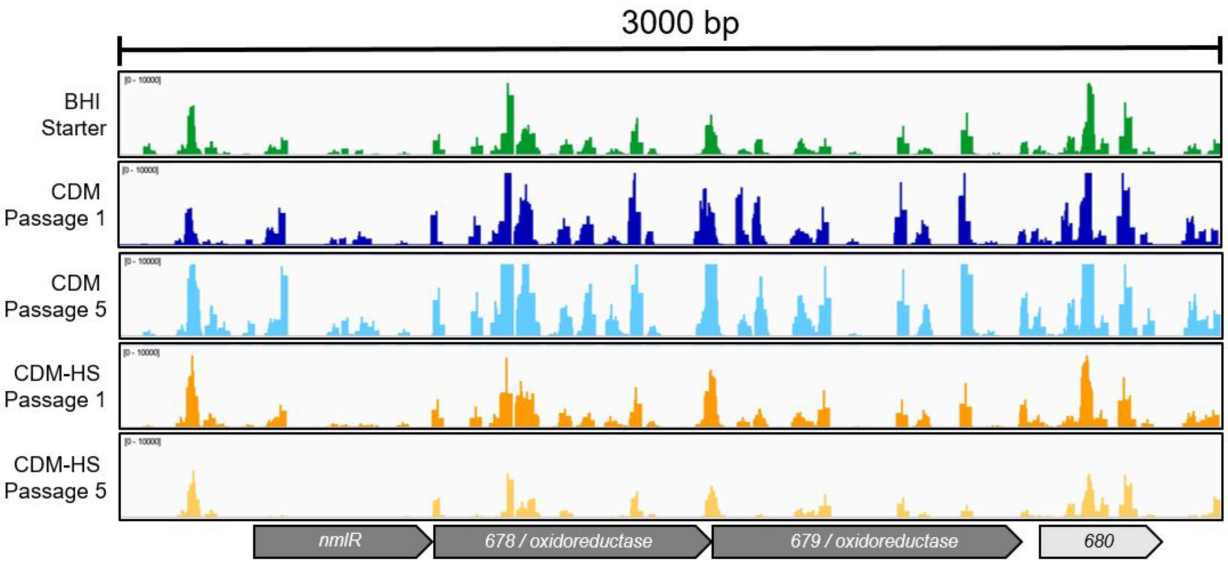
3000 bp
BHI
Starter
CDM
Passage 1
CDM
Passage 5
CDM-HS
Passage 1
CDM-HS
Passage 5
nmlR
678 / oxidoreductase
679 / oxidoreductase
680

D

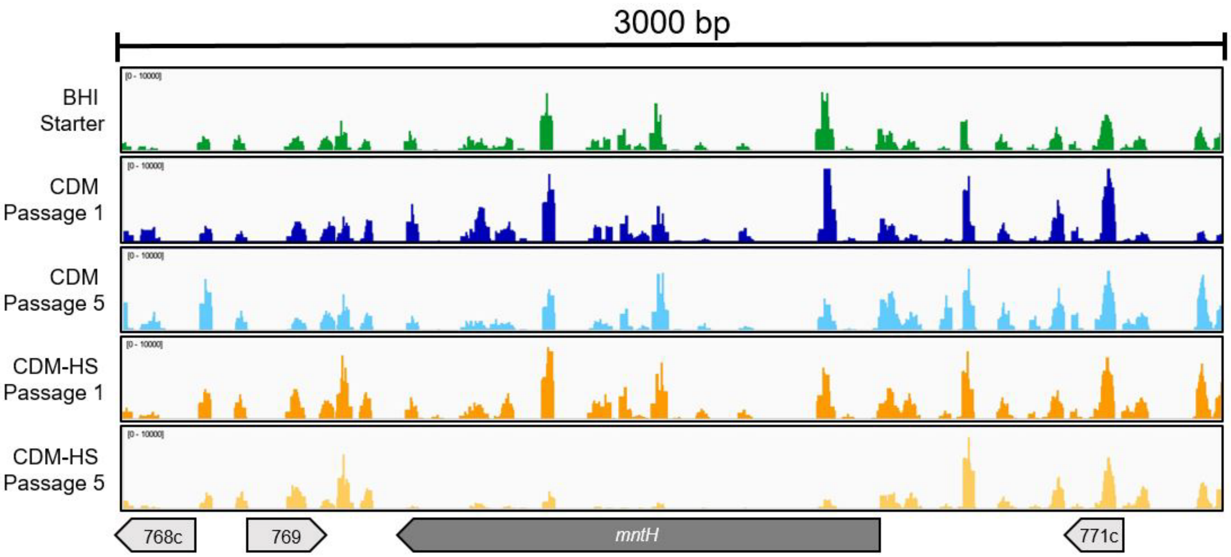
3000 bp
BHI
Starter
CDM
Passage 1
CDM
Passage 5
CDM-HS
Passage 1
CDM-HS
Passage 5
768c
769
mntH
771c

E

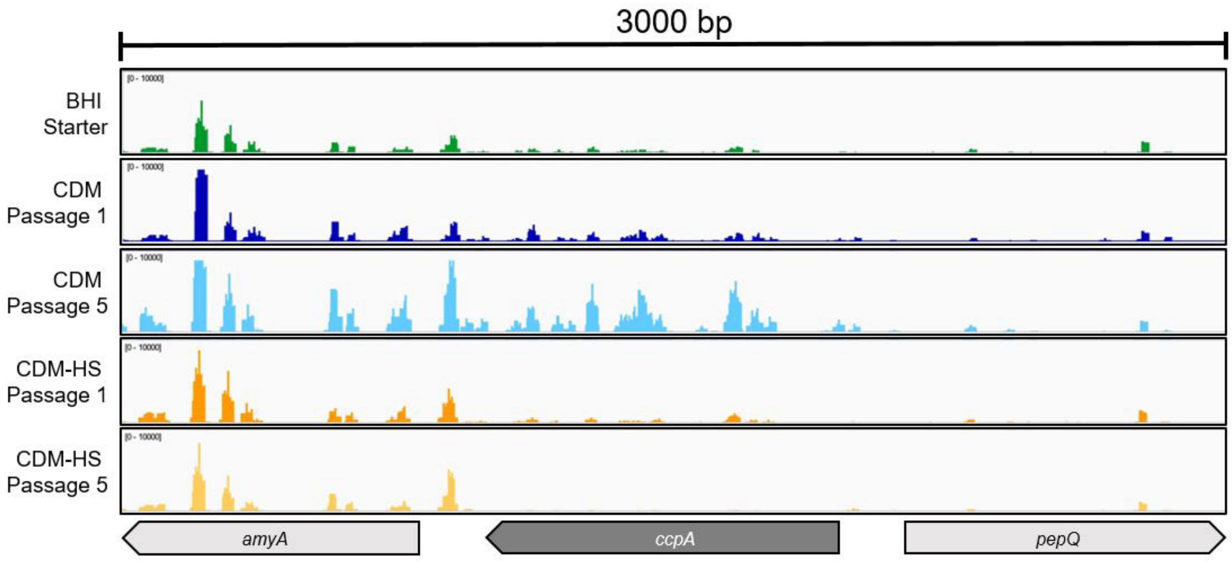

**Supplemental Figure 3.** ***Tn-Seq read coverage across selected genomic loci in Streptococcus mutans.*** Integrative Genomics Viewer (IGV) visualization of Tn-Seq read coverage across (A) the *adhABCD–lplA* locus, (B) *pur* operon, (C) SMU.677 – 679, (D) *mntH*, and (E) *ccpA* in *S. mutans* UA159. Tracks represent the BHI starter culture (green), CDM passage 1 (dark blue), CDM passage 5 (light blue), saliva-supplemented CDM passage 1 (orange), and saliva-supplemented CDM passage 5 (yellow). Gene positions and transcriptional orientations are indicated by arrows below each panel, with dark gray indicating genes with significant depletion in recovered reads in the CDM-HS passage 5 condition. Coverage tracks represent a randomly chosen individual biological replicate and display normalized coverage across tracks. Vertical scales are indicated within each track (top left).

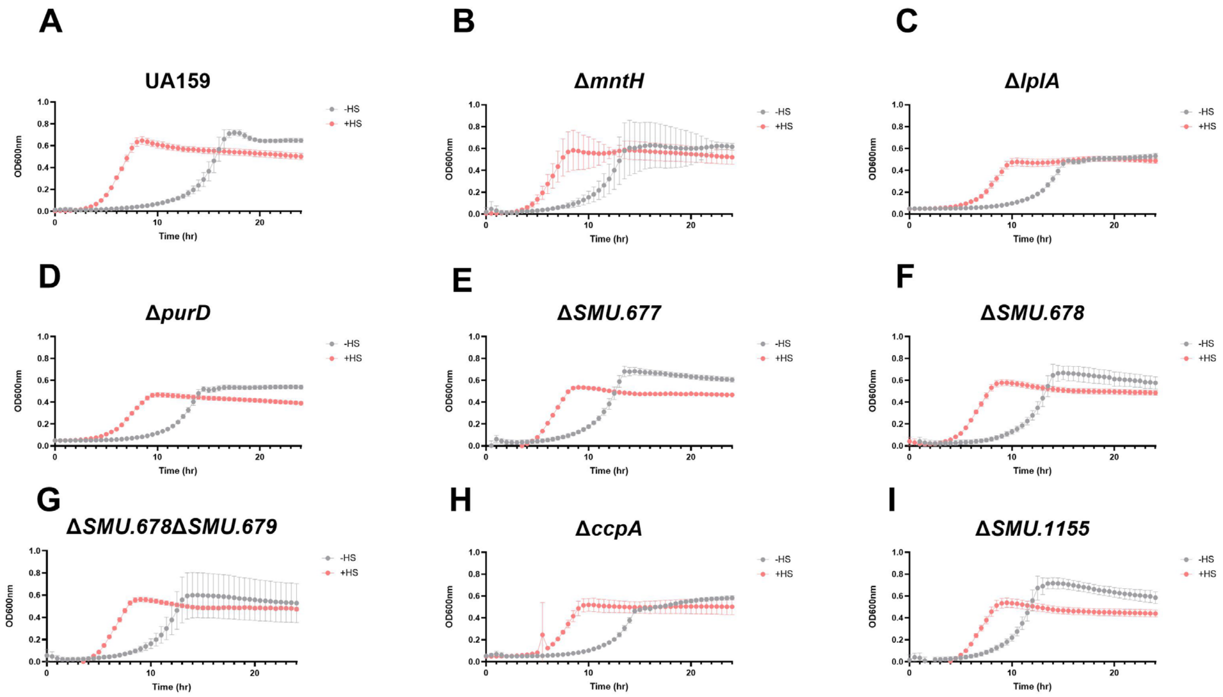

**Supplemental Figure 4.** ***Growth of Streptococcus mutans UA159 and deletion mutants with and without saliva*****.** Growth curves of (A) UA159 (wild-type), (B) Δ*mntH*, (C) Δ*lplA*, (D) Δ*purD*, (E) ΔSMU.677, (F) ΔSMU.678, (G) ΔSMU.678ΔSMU.679, (H) Δ*ccpA*, and (I) ΔSMU.1155 cultured in CDM without saliva (−HS; gray circles) or supplemented with 50% (v/v) pooled human saliva (+HS; red squares). Growth was monitored by optical density at 600 nm every 30 min for 24 hr.

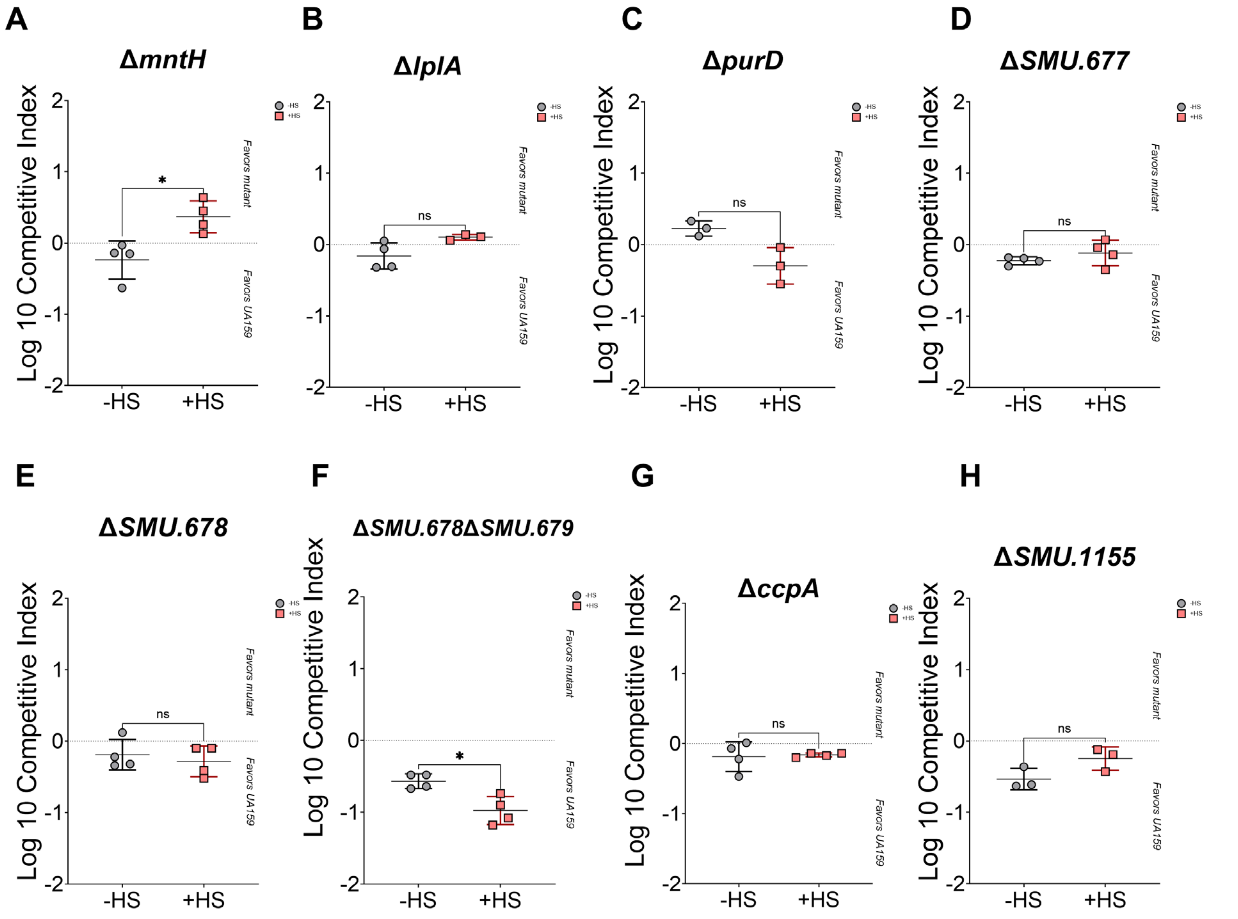

**Supplemental Figure 5.** ***Competitive fitness of Streptococcus mutans deletion mutants against the isogenic parent.*** Competitive indices of (A) Δ*mntH*, (B) Δ*lplA*, (C) Δ*purD*, (D) ΔSMU.677, (E) ΔSMU.678, (F) ΔSMU.678ΔSMU.679, (G) Δ*ccpA*, and (H) ΔSMU.1155 (control strain – not identified by Tn-Seq) during competition with *S. mutans* UA159. Each mutant was inoculated with the parent at a 1:1 ratio in CDM without saliva (−HS; gray circles) or supplemented with 50% (v/v) pooled human saliva (+HS; red squares). Cultures were incubated for 24 hr before plating for CFU enumeration. Positive values favor the mutant, negative values favor UA159, and zero indicates no change in relative abundance. Symbols represent independent competition cultures (n = 4 per condition), with the overall mean indicated by a horizontal line. Differences between −HS and +HS conditions were assessed using unpaired t-tests. *, $P \leq 0.05$; ns, not significant ($P > 0.05$).

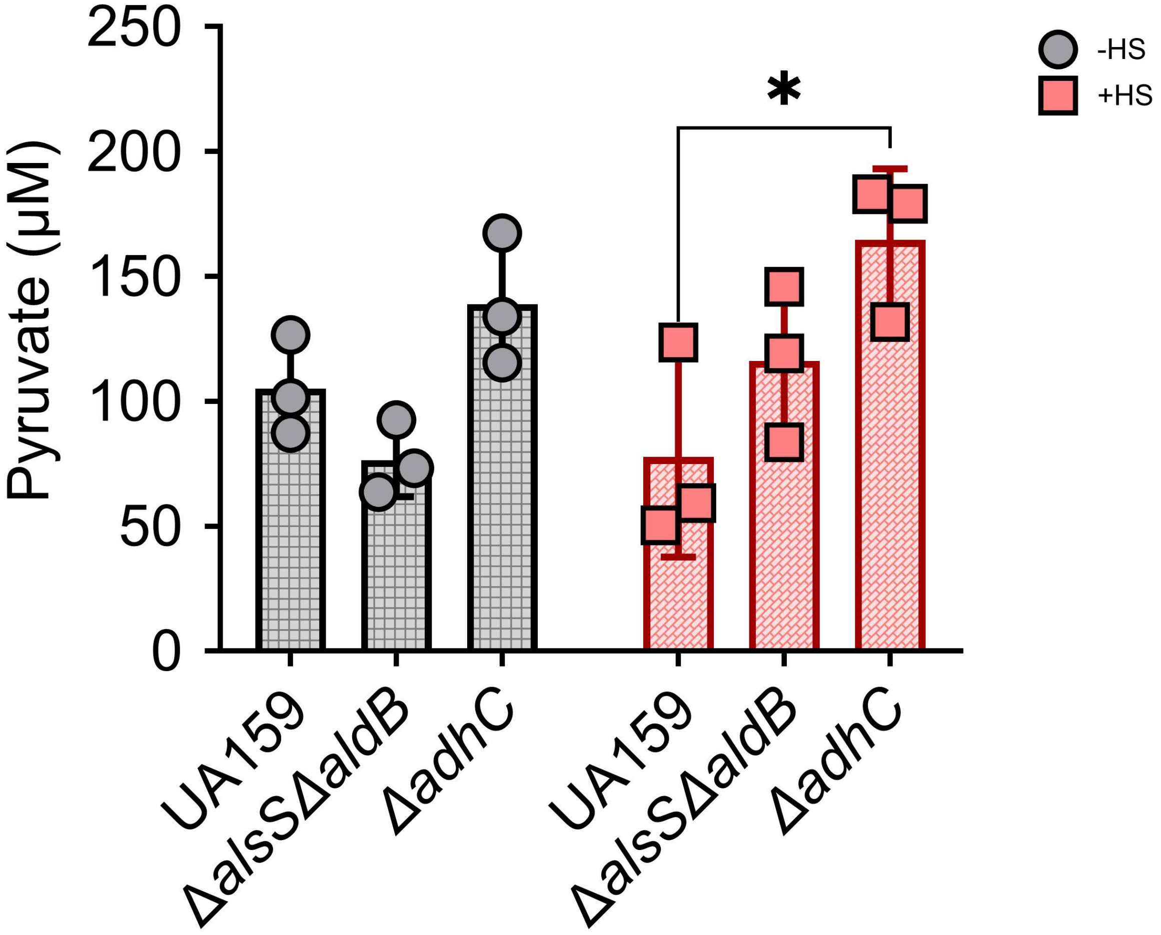

**Supplemental Figure 6.** ***Increased pyruvate levels in whole-cell lysates of saliva-grown Streptococcus mutans lacking adhC.*** Pyruvate concentrations measured in whole-cell lysates of *S. mutans* UA159, Δ*alsS*Δ*aldB*, and Δ*adhC* cultured in CDM without saliva (−HS; gray circles) or supplemented with 50% (v/v) pooled human saliva (+HS; red squares). Symbols represent independent cultures (n = 3 per group). *, P ≤ 0.05.

#### SUPPLEMENTAL TABLES

**Supplemental Table 1. *Differentially expressed genes* (DEGs) of Streptococcus mutans UA159 cultured in saliva by RNA-Seq.*** Red, upregulated genes. Blue, downregulated genes. Table sorted with must upregulated shown first.

*Differentially expressed genes = genes with ≥4 -Log10 *P* value and fold change (FC) ≥ (-)1.5.

| Geneid | Gene | Product | Log2 Fold Change | Fold Change | P value | Log10 P value |
|---|---|---|---|---|---|---|
| SMU_310 | | sorbitol operon activator | 2.252770202 | 4.765971097 | 5.45E-23 | 22.26 |
| SMU_131 | *lplA* | lipoate-protein ligase | 2.234435445 | 4.705785121 | 7.52E-49 | 48.12 |
| SMU_1004 | *gtfB* | glucosyltransferase-I | 2.162810614 | 4.477863695 | 7.66E-29 | 28.12 |
| SMU_130 | *adhD* | dihydrolipoamide dehydrogenase | 2.128540228 | 4.372748058 | 4.74E-35 | 34.32 |
| SMU_308 | | sorbitol-6-phosphate 2-dehydrogenase | 2.092260541 | 4.264156953 | 1.61E-15 | 14.79 |
| SMU_1396 | *gbpC* | glucan-binding protein C | 2.076649774 | 4.218265124 | 1.71E-28 | 27.77 |
| SMU_311 | | PTS system, sorbitol (glucitol) phosphotransferase enzyme IIC2 | 2.076307397 | 4.217264177 | 3.86E-20 | 19.41 |
| SMU_312 | | PTS system, sorbitol phosphotransferase enzyme IIBC | 2.019911435 | 4.055588944 | 5.05E-20 | 19.30 |
| SMU_496 | *cysK* | cysteine synthetase A | 1.935405796 | 3.824856961 | 6.32E-28 | 27.20 |
| SMU_309 | | regulator of sorbitol operon | 1.934918972 | 3.823566517 | 2.82E-16 | 15.55 |
| SMU_313 | | PTS system, sorbitol-specific enzyme IIA | 1.906004152 | 3.747696573 | 3.56E-13 | 12.45 |
| SMU_1397c | | possible transcriptional repressor | 1.755957745 | 3.377504624 | 0.000130733 | 3.88 |
| SMU_314 | | hypothetical protein | 1.750006807 | 3.363601532 | 7.66E-10 | 9.12 |
| SMU_1597c | | conserved hypothetical protein | 1.744268695 | 3.350249861 | 9.05703E-06 | 5.04 |
| SMU_1395c | | hypothetical protein | 1.689427015 | 3.225285818 | 2.91278E-05 | 4.54 |
| SMU_132 | | amino acid amidohydrolase (hippurate amidohydrolase) | 1.662238092 | 3.165071502 | 1.90E-16 | 15.72 |
| SMU_1117 | *naoX* | H2O-forming NADH Oxidase | 1.619446784 | 3.072571929 | 1.14E-20 | 19.94 |
| SMU_1116c | | hypothetical protein | 1.612483378 | 3.057777384 | 1.21E-13 | 12.92 |
| SMU_1596 | *ptcC* | PTS system, cellobiose-specific IIC component | 1.567207632 | 2.963306044 | 6.45E-11 | 10.19 |
| SMU_129 | *adhC* | dihydrolipoamide S-acetyltransferase | 1.566523404 | 2.961900968 | 9.70E-23 | 22.01 |
| SMU_1599 | *celR* | transcriptional regulator, required for cellobiose uptake and metabolism | 1.533011804 | 2.893893449 | 3.14E-09 | 8.50 |
| SMU_128 | *adhB* | acetoin dehydrogenase (TPP-dependent) E1 component beta subunit | 1.484920061 | 2.799016634 | 9.08E-17 | 16.04 |
| SMU_438c | | (R)-2-hydroxyglutaryl-CoA dehydratase activator-related protein | 1.482952374 | 2.795201667 | 6.26E-18 | 17.20 |
| SMU_55 | | hypothetical protein | 1.370508676 | 2.585617157 | 0.000363645 | 3.44 |
| SMU_127 | *adhA* | acetoin dehydrogenase (TPP-dependent) E1 component alpha subunit | 1.231933527 | 2.348815717 | 2.27E-14 | 13.64 |
| SMU_283 | | hypothetical protein | 1.226622215 | 2.340184401 | 0.00022547 | 3.65 |
| SMU_957 | | 50S ribosomal protein L10 | 1.123704594 | 2.179057995 | 9.24905E-05 | 4.03 |
| SMU_459 | | amino acid ABC transporter, substrate-binding protein | 1.061387984 | 2.086938351 | 1.29712E-06 | 5.89 |
| SMU_461 | | amino acid ABC transporter, ATP-binding protein | 1.045864765 | 2.064603531 | 7.36E-07 | 6.13 |
| SMU_1005 | *gtfC* | glucosyltransferase-SI | 0.998831052 | 1.998380151 | 1.12E-07 | 6.95 |
| SMU_1841 | *scrA* | PTS system, sucrose-specific IIABC component | 0.979089023 | 1.971220307 | 8.28E-11 | 10.08 |
| SMU_145 | | major facilitator superfamily transporter, efflux protein | 0.96767983 | 1.955692882 | 2.55264E-05 | 4.59 |

| | | | | | | |
|---|---|---|---|---|---|---|
| SMU_460 | | amino acid ABC transporter, permease | 0.964115972 | 1.950867739 | 5.40107E-06 | 5.27 |
| SMU_1878 | *ptnC* | mannose PTS system component IIC | 0.957118392 | 1.941428252 | 5.81E-09 | 8.24 |
| SMU_435 | | N-acetylglucosamine-6-phosphate deacetylase | 0.948556647 | 1.929940871 | 2.42104E-06 | 5.62 |
| SMU_1591 | *ccpA* | catabolite control protein A | 0.9481072 | 1.929339725 | 2.14E-09 | 8.67 |
| SMU_1928 | *psaB* | protein secretion ABC transport permease | 0.937903166 | 1.915741847 | 3.44E-09 | 8.46 |
| SMU_426 | *copA* | copper-transporting ATPase | 0.932294778 | 1.908308975 | 2.71E-08 | 7.57 |
| SMU_627 | | conserved hypothetical protein | 0.92284991 | 1.895856685 | 0.000376194 | 3.42 |
| SMU_494 | | transaldolase family protein | 0.912286202 | 1.882025534 | 7.14227E-05 | 4.15 |
| SMU_427 | *copZ* | copper chaperone | 0.898679781 | 1.864359115 | 0.000134361 | 3.87 |
| SMU_1590 | *amyA* | intracellular a-amylase | 0.898540351 | 1.864178943 | 4.92E-08 | 7.31 |
| SMU_86 | | conserved hypothetical protein | 0.852676067 | 1.805847501 | 0.000212038 | 3.67 |
| SMU_1734 | *accA* | acetyl-CoA carboxylase alpha subunit | 0.852520022 | 1.805652188 | 7.82465E-06 | 5.11 |
| SMU_271 | *ptxB* | PTS system, enzyme IIB component | 0.848587505 | 1.80073702 | 0.000484869 | 3.31 |
| SMU_277 | | hypothetical protein | 0.838576678 | 1.788285 | 0.0002504 | 3.60 |
| SMU_1737 | *fabZ* | (3R)-hydroxymyristoyl-(acyl carrier protein) dehydratase | 0.834856256 | 1.783679312 | 1.38726E-05 | 4.86 |
| SMU_112c | | transcriptional regulator, RpiR family | 0.831679111 | 1.779755563 | 9.57397E-05 | 4.02 |
| SMU_1425 | *clpB* | ATP-dependent Clp protease, ATP-binding subunit ClpB | 0.825368279 | 1.771987319 | 2.71851E-05 | 4.57 |
| SMU_2038 | *pttB* | phosphotransferase system, trehalose-specific IIBC component (EIIBC-tre) | 0.808439756 | 1.751316407 | 0.000165661 | 3.78 |
| SMU_2037 | *treA* | trehalose-6-phosphate hydrolase | 0.803408438 | 1.745219427 | 0.000328537 | 3.48 |
| SMU_1877 | *ptnA* | mannose PTS system component IIAB | 0.801232487 | 1.742589176 | 3.16878E-06 | 5.50 |
| SMU_2009 | *rs5* | 30S ribosomal protein S5 | 0.78845109 | 1.727219085 | 2.22685E-05 | 4.65 |
| SMU_1879 | | mannose PTS system component IID | 0.760652054 | 1.694256204 | 8.45E-07 | 6.07 |
| SMU_274 | | hexulose-6-phosphate isomerase | 0.758699502 | 1.691964738 | 0.000184924 | 3.73 |
| SMU_2017 | *rl14* | 50S ribosomal protein L14 | 0.75439515 | 1.686924201 | 7.53361E-06 | 5.12 |
| SMU_34 | *purM* | phosphoribosylformylglycinamide cyclo-ligase (AIRS) | 0.744170624 | 1.675011063 | 7.67246E-05 | 4.12 |
| SMU_37 | *purH* | phosphoribosylaminoimidazolecarboxamide formyltransferase / IMP cyclohydrolase | 0.74139907 | 1.671796297 | 0.000109736 | 3.96 |
| SMU_59 | *purB* | adenylosuccinate lyase | 0.728279701 | 1.656662476 | 8.80536E-05 | 4.06 |
| SMU_1947 | *nusG* | transcription antitermination factor | 0.716882719 | 1.643626754 | 0.00050275 | 3.30 |
| SMU_2020 | *rl16* | 50S ribosomal protein L16 | 0.707150244 | 1.632576102 | 2.57844E-05 | 4.59 |
| SMU_493 | *pfl2* | formate acetyltransferase (pyruvate-formate lyase) | 0.703141393 | 1.628045921 | 0.000415528 | 3.38 |
| SMU_36 | | conserved hypothetical protein (eukaryotic-like) | 0.702324959 | 1.627124856 | 0.00029534 | 3.53 |
| SMU_530c | | conserved hypothetical protein | 0.695622678 | 1.619583307 | 2.53508E-05 | 4.60 |
| SMU_35 | *purN* | phosphoribosylglycinamide formyltransferase | 0.67443993 | 1.595977077 | 0.00083157 | 3.08 |
| SMU_1296 | | glutathione S-transferase | 0.672254268 | 1.593561026 | 0.000176453 | 3.75 |
| SMU_2010 | *rl18* | 50S ribosomal protein L18 | 0.669862907 | 1.590921782 | 0.000535806 | 3.27 |
| SMU_51 | *purK* | phosphoribosylaminoimidazole carboxylase, ATPase subunit | 0.660044422 | 1.580131277 | 0.000166021 | 3.78 |
| SMU_888 | *galE* | UDP-glucose 4-epimerase | 0.656586385 | 1.57634835 | 2.25019E-05 | 4.65 |
| SMU_157 | *cysE* | serine O-acetyltransferase | 0.634237788 | 1.552117511 | 0.000705467 | 3.15 |
| SMU_1822 | *gatA* | aspartyl-tRNA synthetase | 0.633835815 | 1.55168511 | 7.30591E-06 | 5.14 |
| SMU_48 | *purD* | phosphoribosylamine-glycine ligase | 0.632988505 | 1.550774056 | 0.000679691 | 3.17 |

| | | | | | | |
|---|---|---|---|---|---|---|
| SMU_1539 | *glgB* | 1,4-alpha-glucan branching enzyme | 0.59504239 | 1.510516968 | 0.00032534 | 3.49 |
| SMU_124 | | transcriptional regulator, MarR family | 0.592439958 | 1.507794649 | 0.000479178 | 3.32 |
| SMU_1434c | | glycosyl transferase | 0.588956777 | 1.504158686 | 3.72787E-05 | 4.43 |
| SMU_1432c | | glycosyl hydrolases family 8 protein; possible beta-glucanase | 0.588067528 | 1.503231838 | 0.000101764 | 3.99 |
| SMU_1735 | *accD* | acetyl-CoA carboxylase beta subunit | 0.586832254 | 1.501945281 | 4.60871E-05 | 4.34 |
| SMU_1451 | *aldB* | alpha-acetolactate decarboxylase | 0.582135872 | 1.497063973 | 0.000679718 | 3.17 |
| SMU_1173 | *cysD* | O-acetylhomoserine sulfhydrylase | -0.606120596 | 1.522160614 | 9.81013E-05 | 4.01 |
| SMU_1412c | | ABC transporter, ATP-binding and permease fusion protein | -0.645388943 | 1.564160919 | 1.33866E-05 | 4.87 |
| SMU_942 | *mvaA* | 3-hydroxy-3-methylglutaryl-CoA reductase | -0.665199273 | 1.585787296 | 0.000111808 | 3.95 |
| SMU_775c | | conserved hypothetical protein | -0.700559511 | 1.625134936 | 8.91561E-06 | 5.05 |
| SMU_1074 | | flavoprotein | -0.710544209 | 1.636421286 | 0.000789239 | 3.10 |
| SMU_746c | | conserved hypothetical protein | -0.719434796 | 1.646536845 | 0.000200979 | 3.70 |
| SMU_1723c | | conserved hypothetical protein | -0.723379551 | 1.651045125 | 0.000261039 | 3.58 |
| SMU_669c | | glutaredoxin | -0.730244669 | 1.658920407 | 0.000686587 | 3.16 |
| SMU_1724c | | rRNA methyltransferase | -0.74174168 | 1.67219336 | 0.00073324 | 3.13 |
| SMU_82 | *dnaK* | chaperone protein, DnaK | -0.76696355 | 1.701684465 | 7.11E-07 | 6.15 |
| SMU_1438c | | conserved hypothetical protein; possible protease | -0.771659408 | 1.707232337 | 0.000132237 | 3.88 |
| SMU_719c | | conserved hypothetical protein | -0.783096486 | 1.720820344 | 3.39222E-05 | 4.47 |
| SMU_104 | | glycosyl hydrolase, alpha-glucosidase | -0.796482 | 1.736860638 | 0.000701845 | 3.15 |
| SMU_102 | | PTS system, IID component | -0.797458147 | 1.73803622 | 0.000440223 | 3.36 |
| SMU_748 | | hypothetical protein | -0.808812186 | 1.751768566 | 7.09344E-05 | 4.15 |
| SMU_837 | | oxidoreductase, aldo/keto reductase family | -0.837126926 | 1.786488871 | 0.000164496 | 3.78 |
| SMU_1995c | | zinc transport transcriptional repressor | -0.926249998 | 1.900330036 | 0.000509254 | 3.29 |
| SMU_1409c | | transcriptional regulator | -0.92913951 | 1.90413994 | 0.000234366 | 3.63 |
| SMU_1068c | | ABC transporter, ATP-binding protein | -0.943078727 | 1.92262677 | 4.39136E-05 | 4.36 |
| SMU_998 | | ABC transporter, ferrichrome-binding protein | -0.972123931 | 1.961726518 | 1.27805E-05 | 4.89 |
| SMU_873 | *metE* | 5-methyltetrahydropteroyltriglutamate--homocysteine S-methyltransferase | -0.988907273 | 1.984681184 | 3.02487E-05 | 4.52 |
| SMU_1097c | | transcriptional regulator | -0.989482695 | 1.985472936 | 0.000361148 | 3.44 |
| SMU_2080 | | conserved hypothetical protein | -0.994660882 | 1.99261209 | 3.53555E-05 | 4.45 |
| SMU_769 | | conserved hypothetical protein | -1.016530287 | 2.023047631 | 3.78585E-06 | 5.42 |
| SMU_997 | | inorganic ion ABC transporter,ATP-binding protein; possible ferrichrome transport system | -1.023250667 | 2.032493399 | 5.53E-07 | 6.26 |
| SMU_1175 | | sodium:alanine (or glycine) symporter | -1.06160089 | 2.087246356 | 2.65913E-06 | 5.58 |
| SMU_874 | | 5-methyltetrahydrofolate--homocysteine methyltransferase | -1.064557029 | 2.091527589 | 2.2943E-06 | 5.64 |
| SMU_1475c | | conserved hypothetical protein | -1.074966528 | 2.106673194 | 6.5136E-05 | 4.19 |
| SMU_149 | | transposase fragment (IS605/IS200-like) | -1.086027243 | 2.122886505 | 0.000442197 | 3.35 |
| SMU_1675 | *metB* | cystathionine gamma-synthase | -1.139689141 | 2.203335426 | 3.57E-07 | 6.45 |
| SMU_1674 | *metC* | aminotransferase class II | -1.141748915 | 2.206483434 | 2.37E-07 | 6.63 |
| SMU_1521 | | amino acid (glutamine) ABC transporter permease | -1.144636226 | 2.210903759 | 1.22E-12 | 11.92 |
| SMU_768c | | conserved hypothetical protein | -1.220066172 | 2.32957402 | 3.31857E-05 | 4.48 |
| SMU_1302 | *adcA* | zinc ABC transporter, substrate-binding protein | -1.247348182 | 2.374046481 | 1.62E-07 | 6.79 |

| | | | | | | |
|---|---|---|---|---|---|---|
| SMU_770c | | manganese transporter/possible HitA ferric iron-binding periplasmic protein | -1.258920656 | 2.393166305 | 7.87E-14 | 13.10 |
| SMU_913 | | NADP-specific glutamate dehydrogenase | -1.275354084 | 2.420582195 | 3.54E-14 | 13.45 |
| SMU_1519 | *glnQ* | glutamine ABC transport, ATP-binding protein | -1.288434549 | 2.442628649 | 1.06E-13 | 12.98 |
| SMU_1522 | *glnP* | amino acid (glutamine) ABC transporter permease | -1.373001795 | 2.590089221 | 7.50E-19 | 18.12 |
| SMU_805c | | amino acid ABC transporter, ATP-binding protein | -1.387029585 | 2.615396329 | 3.17E-12 | 11.50 |
| SMU_806c | | glutamine ABC transporter, permease and solute binding protein | -1.38862596 | 2.618291927 | 9.19E-16 | 15.04 |
| SMU_2084c | | conserved hypothetical protein (possible arsenate reductase) | -1.392806904 | 2.625890768 | 2.80E-17 | 16.55 |
| SMU_367 | | Streptococcus-specific protein; similar to glucan-binding protein | -1.402306467 | 2.643238247 | 2.83E-15 | 14.55 |
| SMU_195c | | hypothetical protein | -1.438132245 | 2.709698328 | 9.04344E-05 | 4.04 |
| SMU_1520 | | amino acid ABC transporter, substrate-binding protein | -1.479356874 | 2.788244109 | 9.52E-15 | 14.02 |
| SMU_2027 | | transcriptional regulator/repressor | -1.516605178 | 2.861169917 | 2.05E-11 | 10.69 |
| SMU_1750c | | hypothetical protein | -1.544260052 | 2.91654443 | 8.13E-08 | 7.09 |
| SMU_1487 | | conserved hypothetical protein | -1.549833615 | 2.927833707 | 2.72E-09 | 8.57 |
| SMU_186 | *sloR* | metal-dependent transcriptional regulator (probable DtxR homolog) | -1.627731708 | 3.090267461 | 4.20E-12 | 11.38 |
| SMU_148 | *adhE* | alcohol-acetaldehyde dehydrogenase | -1.666571152 | 3.17459192 | 1.56E-12 | 11.81 |
| SMU_197c | | hypothetical protein | -1.802226318 | 3.487580019 | 4.49588E-05 | 4.35 |
| SMU_202c | | conserved hypothetical protein/Streptococcus-specific protein | -1.806288477 | 3.497413745 | 0.000715317 | 3.15 |
| SMU_193c | | conserved hypothetical protein | -1.826041488 | 3.545628759 | 0.000518386 | 3.29 |
| SMU_214c | | hypothetical protein | -1.828317032 | 3.551225646 | 0.000757237 | 3.12 |
| SMU_198c | | conjugative transposon protein | -1.885343572 | 3.694408946 | 1.82372E-06 | 5.74 |
| SMU_196c | | immunogenic secreted protein (transfer protein) | -1.892031472 | 3.711574872 | 2.74E-13 | 12.56 |
| SMU_1754c | | conserved hypothetical protein | -2.006489699 | 4.018033798 | 4.76E-16 | 15.32 |
| SMU_211c | | hypothetical protein | -2.030334544 | 4.084995656 | 5.16505E-05 | 4.29 |
| SMU_183 | *sloB* | manganese ABC transporter permease element | -2.031142681 | 4.087284539 | 6.76E-12 | 11.17 |
| SMU_1760c | | conserved hypothetical protein | -2.065512825 | 4.185827402 | 2.45E-16 | 15.61 |
| SMU_210c | | hypothetical protein | -2.076373716 | 4.217458043 | 0.000283004 | 3.55 |
| SMU_366 | *gltB* | glutamate synthase, small subunit | -2.09459345 | 4.271057883 | 8.08E-45 | 44.09 |
| SMU_1757c | | conserved hypothetical protein | -2.113478019 | 4.327332624 | 4.29E-14 | 13.37 |
| SMU_1752c | | hypothetical protein | -2.1198401 | 4.346457688 | 7.86E-22 | 21.10 |
| SMU_365 | *gltA* | glutamate synthase, large subunit | -2.207919434 | 4.620085128 | 2.31E-44 | 43.64 |
| SMU_200c | | hypothetical protein | -2.242244461 | 4.731325645 | 0.000302112 | 3.52 |
| SMU_1753c | | conserved hypothetical protein | -2.27120435 | 4.827259386 | 1.54E-23 | 22.81 |
| SMU_182 | *sloA* | ABC transporter, ATP-binding protein, iron and/or manganese | -2.274810693 | 4.839341309 | 1.57E-14 | 13.80 |
| SMU_208c | | conserved hypothetical protein, FtsK/SpoIIIE family | -2.315853769 | 4.978992246 | 2.01E-07 | 6.70 |
| SMU_1758c | | conserved hypothetical protein | -2.321877176 | 4.999823533 | 4.64E-29 | 28.33 |
| SMU_1762c | | conserved hypothetical protein | -2.38057591 | 5.207445768 | 4.00E-20 | 19.40 |
| SMU_184 | *sloC* | ABC transporter element, iron predicted binding protein | -2.430460268 | 5.39065383 | 6.80E-18 | 17.17 |
| SMU_1755c | | conserved hypothetical protein | -2.431860504 | 5.395888378 | 3.18E-24 | 23.50 |
| SMU_207c | | transcriptional regulator | -2.441304591 | 5.431326496 | 8.50E-10 | 9.07 |
| SMU_1187 | *glmS* | L-glutamine-D-fructose-6-phosphate amidotransferase | -2.490790502 | 5.62085852 | 1.17E-48 | 47.93 |

| | | | | | | |
|---|---|---|---|---|---|---|
| SMU_1761c | | conserved hypothetical protein | -2.531640024 | 5.782286223 | 1.39E-22 | 21.86 |
| SMU_363 | *glnR* | transcriptional regulator; glutamine synthetase repressor | -2.650880493 | 6.280504677 | 6.47E-59 | 58.19 |
| SMU_1763c | | conserved hypothetical protein | -2.666725669 | 6.349863893 | 9.28E-26 | 25.03 |
| SMU_185 | | hypothetical protein | -2.684685031 | 6.429404106 | 7.75E-15 | 14.11 |
| SMU_205c | | conserved hypothetical protein | -2.818231014 | 7.052970543 | 4.49E-08 | 7.35 |
| SMU_209c | | hypothetical protein | -2.866374581 | 7.292303369 | 1.97E-09 | 8.71 |
| SMU_1764c | | conserved hypothetical protein | -2.887482133 | 7.399778736 | 5.13E-28 | 27.29 |
| SMU_364 | *glnA* | glutamine synthetase type 1 | -2.932231098 | 7.632898983 | 4.73E-44 | 43.33 |
| SMU_673 | | conserved hypothetical protein | -3.805718608 | 13.98413011 | 1.58E-76 | 75.80 |
| SMU_1657c | | nitrogen regulatory protein PII | -5.279218089 | 38.8331838 | 2.15E-70 | 69.67 |
| SMU_1658 | *nrgA* | ammonium transporter, NrgA protein | -5.350790147 | 40.80828416 | 2.74E-90 | 89.56 |
| SMU_672 | *idh* | isocitrate dehydrogenase | -5.463532209 | 44.12523966 | 6.33E-186 | 185.20 |
| SMU_670 | *citB* | aconitate hydratase; aconitase A | -5.62518219 | 49.35697903 | 3.37E-175 | 174.47 |
| SMU_671 | *citZ* | citrate synthase | -5.671569645 | 50.96975966 | 2.58E-181 | 180.59 |

**Supplemental Table 2. *Significant* under- and over-represented genes in CDM-Saliva by Tn-Seq at Passage 5.*** Genes that were found to be over-represented in CDM-Saliva passage 5 (green) or under-represented (red).

*Significiant = genes with ≥4 Log10 *P* value and CDM_HS p5 fold change (FC) ≥ (-)2.

| Gene ID | Gene | Product | CDM_HSp5 Log2 FC | CDM_HS p5 FC | P value | Log 10 P value |
|---|---|---|---|---|---|---|
| **SMU_1691** | *dltA* | D-alanine-D-alanyl carrier protein ligase | 15.41 | 43577.34 | 0.00 | **5.72** |
| **SMU_1690** | *dltB* | integral membrane protein, D-alanyl transfer | 14.21 | 18940.91 | 0.00 | **5.03** |
| **SMU_831** | | conserved hypothetical protein | 13.65 | 12845.14 | 0.00 | **4.15** |
| **SMU_747c** | | conserved hypothetical protein, permease-like | 13.30 | 10074.95 | 0.00 | **4.43** |
| **SMU_1689** | *dltC* | D-alanyl carrier protein | 13.26 | 9832.52 | 0.00 | **4.95** |
| **SMU_746c** | | conserved hypothetical protein | 12.97 | 8012.15 | 0.00 | **4.54** |
| **SMU_1080c** | | conserved hypothetical protein | 12.94 | 7860.16 | 0.00 | **7.96** |
| **SMU_547** | | conserved hypothetical protein | 12.47 | 5675.30 | 0.00 | **4.92** |
| **SMU_806c** | | glutamine ABC transporter, permease and solute binding protein | 12.36 | 5270.84 | 0.00 | **12.90** |
| **SMU_454** | *ftsL* | cell division protein | 11.07 | 2157.25 | 0.00 | **6.02** |
| **SMU_1100c** | | permease | 10.83 | 1817.10 | 0.00 | **5.59** |
| **SMU_1688** | *dltD* | integral membrane protein | 9.81 | 895.75 | 0.00 | **5.44** |
| **SMU_758c** | | conserved hypothetical protein | 9.61 | 782.43 | 0.00 | **6.85** |
| **SMU_786** | *pheA* | prephenate dehydratase | 9.02 | 518.87 | 0.00 | **7.24** |
| **SMU_483** | | phosphoprotein phosphatase (pppL protein) | 8.94 | 492.05 | 0.00 | **4.02** |
| **SMU_948** | | conserved hypothetical protein | 8.14 | 282.39 | 0.00 | **6.37** |
| **SMU_1428c** | | conserved hypothetical protein | 8.03 | 261.21 | 0.00 | **5.89** |
| **SMU_1361c** | | transcriptional regulator | 7.73 | 212.64 | 0.00 | **4.71** |
| **SMU_560c** | | conserved hypothetical protein | 7.57 | 189.89 | 0.00 | **6.62** |
| **SMU_1061** | *ylxM* | DNA-binding protein | 7.48 | 178.97 | 0.00 | **8.15** |
| **SMU_73** | | conserved hypothetical protein | 7.27 | 154.47 | 0.00 | **8.96** |
| **SMU_1833** | *recG* | ATP-dependent DNA helicase | 7.22 | 149.52 | 0.00 | **4.24** |
| **SMU_757** | | conserved hypothetical protein | 6.83 | 113.77 | 0.00 | **6.09** |
| **SMU_554** | *ylmF* | conserved hypothetical protein | 6.81 | 111.93 | 0.00 | **4.90** |
| **SMU_1745c** | | transcriptional regulator, MarR family | 6.36 | 82.27 | 0.00 | **5.03** |
| **SMU_72** | | conserved hypothetical protein | 6.15 | 70.88 | 0.00 | **6.37** |
| **SMU_835** | | conserved hypothetical protein (possible membrane protein) | 6.10 | 68.44 | 0.00 | **5.61** |
| **SMU_1529** | *atpC* | ATPase, gamma subunit | 5.99 | 63.76 | 0.00 | **4.94** |
| **SMU_1649** | *exoA* | exodeoxyribonuclease III/ Smx nuclease | 5.96 | 62.45 | 0.00 | **4.98** |
| **SMU_921** | | transcriptional regulator | 5.91 | 59.94 | 0.00 | **4.37** |
| **SMU_245** | *mecA* | negative regulator of genetic competence | 5.77 | 54.75 | 0.00 | **4.61** |
| **SMU_744** | *ftsY* | cell division protein FtsY; signal recognition particle (docking protein) | 5.45 | 43.78 | 0.00 | **4.19** |
| **SMU_1975c** | | conserved hypothetical protein | 5.35 | 40.86 | 0.00 | **6.16** |
| **SMU_692** | | possible pore-forming peptide | 5.30 | 39.42 | 0.00 | **4.25** |
| **SMU_1741** | *fabD* | [acyl-carrier-protein] S-malonyltransferase | 5.27 | 38.68 | 0.00 | **5.00** |

| | | | | | | |
|---|---|---|---|---|---|---|
| **SMU_582** | | farnesyl diphosphate synthase (geranyltranstransferase) | 5.17 | 35.94 | 0.00 | **6.88** |
| **SMU_1976c** | | conserved hypothetical protein | 5.14 | 35.21 | 0.00 | **11.17** |
| **SMU_937** | | diphosphomevalonate decarboxylase | 5.10 | 34.30 | 0.00 | **4.14** |
| **SMU_717** | *murM* | beta-lactam resistance factor murM; Serine / alanine adding enzyme | 4.99 | 31.67 | 0.00 | **6.08** |
| **SMU_2084c** | | conserved hypothetical protein (possible arsenate reductase) | 4.97 | 31.36 | 0.00 | **7.09** |
| **SMU_550** | *divIB* | cell division protein | 4.93 | 30.44 | 0.00 | **6.07** |
| **SMU_805c** | | amino acid ABC transporter, ATP-binding protein | 4.88 | 29.54 | 0.00 | **9.36** |
| **SMU_557** | *divIVA* | cell division initiation protein | 4.71 | 26.10 | 0.00 | **5.88** |
| **SMU_91** | *ropA* | peptidyl-prolyl isomerase, trigger factor | 4.45 | 21.93 | 0.00 | **7.01** |
| **SMU_1949** | *pbp2a* | penicillin-binding protein 2a | 4.38 | 20.84 | 0.00 | **4.17** |
| **SMU_410** | *brpA* | transcriptional regulator | 4.29 | 19.62 | 0.00 | **5.44** |
| **SMU_1270** | *hisD* | histidinol dehydrogenase | 4.08 | 16.97 | 0.00 | **4.76** |
| **SMU_148** | *adhE* | alcohol-acetaldehyde dehydrogenase | 3.99 | 15.90 | 0.00 | **9.16** |
| **SMU_1878** | *ptnC* | mannose PTS system component IIC | 3.91 | 15.05 | 0.00 | **4.58** |
| **SMU_1278c** | | phosphoglycolate phosphatase | 3.76 | 13.58 | 0.00 | **6.13** |
| **SMU_1883** | | conserved hypothetical protein | 3.62 | 12.26 | 0.00 | **5.09** |
| **SMU_707c** | | endolysin | 3.59 | 12.05 | 0.00 | **5.07** |
| **SMU_1427c** | | conserved hypothetical protein | 3.49 | 11.27 | 0.00 | **5.26** |
| **SMU_1064c** | | transcriptional regulator | 3.41 | 10.66 | 0.00 | **6.10** |
| **SMU_1112c** | | conserved hypothetical protein | 3.29 | 9.76 | 0.00 | **5.51** |
| **SMU_486** | | two-component sensor histidine kinase | 3.29 | 9.76 | 0.00 | **7.80** |
| **SMU_703c** | | conserved hypothetical protein | 3.20 | 9.20 | 0.00 | **4.59** |
| **SMU_1533** | *atpG* | ATP synthase subunit A | 3.09 | 8.51 | 0.00 | **4.84** |
| **SMU_66** | | conserved hypothetical protein; possible phosphatidylinositol-4-phosphate 5-kinase | 3.06 | 8.34 | 0.00 | **5.64** |
| **SMU_1876** | | conserved hypothetical protein | 3.05 | 8.28 | 0.00 | **5.00** |
| **SMU_890** | | conserved hypothetical protein | 3.01 | 8.03 | 0.00 | **6.50** |
| **SMU_753** | | conserved hypothetical protein | 3.00 | 8.02 | 0.00 | **5.27** |
| **SMU_1370c** | | | 2.94 | 7.70 | 0.00 | **5.33** |
| **SMU_1268** | *hisB* | imidazoleglycerol-phosphate dehydratase | 2.91 | 7.50 | 0.00 | **6.10** |
| **SMU_485** | | conserved hypothetical protein | 2.91 | 7.49 | 0.00 | **6.08** |
| **SMU_470** | | conserved hypothetical protein | 2.90 | 7.46 | 0.00 | **7.71** |
| **SMU_1113** | *srtA* | sortase | 2.81 | 7.01 | 0.00 | **7.12** |
| **SMU_764** | *ahpC* | alkyl hydroperoxide reductase, subunit C | 2.76 | 6.78 | 0.00 | **5.05** |
| **SMU_402** | *pfl* | pyruvate formate-lyase | 2.55 | 5.85 | 0.00 | **7.43** |
| **SMU_1266** | *hisH* | amidotransferase; possible imidazoleglycerol-phosphate synthase | 2.54 | 5.81 | 0.00 | **5.19** |
| **SMU_354** | | conserved hypothetical protein | 2.50 | 5.66 | 0.00 | **6.98** |
| **SMU_1450** | | amino acid permease | 2.46 | 5.49 | 0.00 | **4.76** |
| **SMU_76** | | N-acetyl-muramidase (N-acetylmuramoyl-L-alanine amidase) | 2.45 | 5.48 | 0.00 | **6.49** |
| **SMU_765** | | alkyl hydroperoxide reductase, subunit F | 2.41 | 5.30 | 0.00 | **5.77** |
| **SMU_337** | | inner membrane protein | 2.37 | 5.17 | 0.00 | **5.16** |
| **SMU_408** | | xanthine/uracil permease family protein | 2.29 | 4.90 | 0.00 | **4.90** |

| | | | | | | |
|---|---|---|---|---|---|---|
| **SMU_1272** | *hisZ* | histidyl-tRNA synthetase | 2.29 | 4.88 | 0.00 | **4.46** |
| **SMU_243** | | conserved hypothetical protein | 2.20 | 4.61 | 0.00 | **8.94** |
| **SMU_1611c** | | multi-drug resistance efflux pump, major facilitator superfamily | 2.18 | 4.54 | 0.00 | **5.60** |
| **SMU_1733c** | | SNF helicase | 2.13 | 4.37 | 0.00 | **5.28** |
| **SMU_67** | | acyltransferase | 2.09 | 4.25 | 0.00 | **5.37** |
| **SMU_1939c** | | amino acid ABC transport ATP-binding protein | 2.03 | 4.10 | 0.00 | **4.04** |
| **SMU_661** | | transcriptional regulator | 2.02 | 4.04 | 0.00 | **5.29** |
| **SMU_242c** | | glutamine ABC transporter, solute binding protein | 2.02 | 4.04 | 0.00 | **5.52** |
| **SMU_1079c** | | ABC transporter, ATP-binding / permease protein | 1.95 | 3.87 | 0.00 | **4.57** |
| **SMU_356** | *purR* | purine operon repressor/xanthine phosphoribosyltransferase | 1.90 | 3.74 | 0.00 | **4.06** |
| **SMU_940c** | | hemolysin III-related protein | 1.89 | 3.70 | 0.00 | **4.47** |
| **SMU_1078c** | | multidrug ABC transporter, ATP-binding and permease protein | 1.87 | 3.67 | 0.00 | **5.64** |
| **SMU_2067** | *csbB* | stress response protein (glycosyltransferase) (sugar transferase) | 1.84 | 3.59 | 0.00 | **5.01** |
| **SMU_648** | *prtM* | protease maturation protein; possible peptidylprolyl isomerase | 1.77 | 3.41 | 0.00 | **5.20** |
| **SMU_1043c** | | phosphate acetyltransferase | 1.75 | 3.37 | 0.00 | **4.04** |
| **SMU_1692** | *pflA* | pyruvate-formate lyase activating enzyme | 1.74 | 3.34 | 0.00 | **5.70** |
| **SMU_412c** | | Hit-like protein | 1.72 | 3.29 | 0.00 | **5.32** |
| **SMU_1661c** | | signal peptidase II | 1.70 | 3.24 | 0.00 | **4.73** |
| **SMU_1118c** | | ribonucleoside ABC transporter permease | 1.68 | 3.21 | 0.00 | **4.88** |
| **SMU_917c** | | 6-pyruvoyl tetrahydropterin synthase, PTPS | 1.67 | 3.18 | 0.00 | **4.30** |
| **SMU_104** | | glycosyl hydrolase, alpha-glucosidase | 1.64 | 3.12 | 0.00 | **5.35** |
| **SMU_105** | | SCR operon transcriptional repressor | 1.62 | 3.07 | 0.00 | **4.26** |
| **SMU_1053** | | possible redox-sensing transcriptional repressor Rex | 1.50 | 2.83 | 0.00 | **4.26** |
| **SMU_2066c** | | conserved hypothetical protein (possible integral membrane protein, ABC transporter) | 1.46 | 2.75 | 0.00 | **5.26** |
| **SMU_366** | *gltB* | glutamate synthase, small subunit | 1.43 | 2.70 | 0.00 | **5.79** |
| **SMU_1674** | *metC* | aminotransferase class II | 1.43 | 2.69 | 0.00 | **5.57** |
| **SMU_701c** | | integral membrane protein | 1.42 | 2.68 | 0.00 | **4.88** |
| **SMU_159** | | conserved hypothetical protein | 1.39 | 2.63 | 0.00 | **4.01** |
| **SMU_365** | *gltA* | glutamate synthase, large subunit | 1.38 | 2.60 | 0.00 | **5.27** |
| **SMU_1452** | *alsS* | alpha-acetolactate synthase | 1.29 | 2.44 | 0.00 | **5.03** |
| **SMU_267c** | | glutamate--cysteine ligase (gamma ECS) | 1.28 | 2.43 | 0.00 | **5.35** |
| **SMU_355** | | CMP-binding factor 1 | 1.24 | 2.36 | 0.00 | **4.33** |
| **SMU_1119c** | | ribonucleoside ABC transporter permease | 1.18 | 2.27 | 0.00 | **4.25** |
| **SMU_1523** | *endA* | competence associated membrane nuclease (DNA-entry nuclease) | 1.15 | 2.21 | 0.00 | **4.45** |
| **SMU_702c** | | transcriptional regulator | 1.13 | 2.19 | 0.00 | **4.02** |
| **SMU_524** | | ABC transporter ATP-binding / permease protein | 1.06 | 2.09 | 0.00 | **5.32** |
| **SMU_235** | | conserved hypothetical protein, SPFH domain/band 7 family | 1.03 | 2.05 | 0.00 | **4.43** |
| **SMU_367** | | Streptococcus-specific protein; similar to glucan-binding protein | 1.01 | 2.02 | 0.00 | **4.04** |
| **SMU_841** | | aminotransferase, class V; possible cysteine desulphurase | -1.12 | -2.18 | 0.00 | **5.33** |
| **SMU_1388** | | ATP-dependent RNA helicase, DEAD/DEAH family | -1.35 | -2.54 | 0.00 | **4.14** |

| | | | | | | |
|---|---|---|---|---|---|---|
| **SMU_1539** | *glgB* | 1,4-alpha-glucan branching enzyme | -1.45 | -2.72 | 0.00 | **4.03** |
| **SMU_1055** | *radC* | DNA repair protein RadC | -1.47 | -2.76 | 0.00 | **4.09** |
| **SMU_872** | | fructose-specific PTS system enzyme IIBC component | -1.48 | -2.80 | 0.00 | **4.40** |
| **SMU_1107c** | | conserved hypothetical protein | -1.50 | -2.83 | 0.00 | **6.21** |
| **SMU_71** | | MATE efflux family protein | -1.55 | -2.93 | 0.00 | **4.65** |
| **SMU_2075c** | | conserved hypothetical protein (possible competence-induced protein) | -1.59 | -3.01 | 0.00 | **5.87** |
| **SMU_1062** | *opuAb* | glycine-betaine binding ABC transporter permease | -1.60 | -3.04 | 0.00 | **5.46** |
| **SMU_770c** | | manganese transporter/possible HitA ferric iron-binding periplasmic protein | -1.61 | -3.06 | 0.00 | **6.19** |
| **SMU_859** | *pyrA* | carbamoyl-phosphate synthase,small subunit | -1.69 | -3.23 | 0.00 | **5.97** |
| **SMU_681** | | conserved hypothetical protein | -1.74 | -3.34 | 0.00 | **5.75** |
| **SMU_679** | | oxidoreductase, aldo/keto reductase family | -1.75 | -3.36 | 0.00 | **6.13** |
| **SMU_678** | | oxidoreductase, aldo/keto reductase family | -1.84 | -3.59 | 0.00 | **4.07** |
| **SMU_860** | *pyrAB* | carbamoyl-phosphate synthase, large subunit | -1.87 | -3.66 | 0.00 | **4.50** |
| **SMU_2146c** | | conserved hypothetical protein | -2.00 | -3.99 | 0.00 | **5.53** |
| **SMU_1711** | *rluB* | ribosomal large subunit pseudouridine synthase B | -2.24 | -4.73 | 0.00 | **5.01** |
| **SMU_834** | | glycosyltransferase | -2.58 | -5.96 | 0.00 | **4.12** |
| **SMU_836** | | hypothetical protein | -2.61 | -6.10 | 0.00 | **5.81** |
| **SMU_870** | | lactose phosphotransferase system repressor/transcriptional repressor of the fructose operon | -2.77 | -6.84 | 0.00 | **5.52** |
| **SMU_128** | *adhB* | acetoin dehydrogenase (TPP-dependent) E1 component beta subunit | -2.88 | -7.37 | 0.00 | **9.72** |
| **SMU_127** | *adhA* | acetoin dehydrogenase (TPP-dependent) E1 component alpha subunit | -2.89 | -7.39 | 0.00 | **7.70** |
| **SMU_774** | | hydrolase, haloacid dehalogenase-like | -2.91 | -7.54 | 0.00 | **6.40** |
| **SMU_721** | | conserved hypothetical protein | -2.97 | -7.83 | 0.00 | **7.21** |
| **SMU_1662** | *holB* | DNA polymerase III delta prime subunit | -3.06 | -8.34 | 0.00 | **4.91** |
| **SMU_871** | *pfkB* | fructose-1-phosphate kinase | -3.10 | -8.55 | 0.00 | **5.11** |
| **SMU_720** | | conserved hypothetical protein | -3.11 | -8.60 | 0.00 | **7.48** |
| **SMU_131** | *lplA* | lipoate-protein ligase | -3.25 | -9.51 | 0.00 | **8.19** |
| **SMU_130** | *adhD* | dihydrolipoamide dehydrogenase | -3.28 | -9.70 | 0.00 | **7.68** |
| **SMU_129** | *adhC* | dihydrolipoamide S-acetyltransferase | -3.44 | -10.88 | 0.00 | **6.75** |
| **SMU_06** | | uncharacterized GTP-binding protein | -3.79 | -13.86 | 0.00 | **5.55** |
| **SMU_677** | | transcriptional regulator, MerR family | -4.05 | -16.59 | 0.00 | **7.84** |
| **SMU_1444c** | | metallo-beta-lactamase superfamily proteinhetical protein | -4.54 | -23.28 | 0.00 | **4.11** |
| **SMU_1591** | *ccpA* | catabolite control protein A | -4.57 | -23.70 | 0.00 | **5.59** |
| **SMU_34** | *purM* | phosphoribosylformylglycinamide cyclo-ligase (AIRS) | -4.68 | -25.68 | 0.00 | **7.70** |
| **SMU_48** | *purD* | phosphoribosylamine-glycine ligase | -4.95 | -30.93 | 0.00 | **5.06** |
| **SMU_32** | *purF* | amidophosphoribosyltransferase | -5.18 | -36.26 | 0.00 | **7.47** |
| **SMU_30** | *purL* | phosphoribosylformylglycinamidine synthase | -5.75 | -53.92 | 0.00 | **10.59** |
| **SMU_35** | *purN* | phosphoribosylglycinamide formyltransferase | -5.94 | -61.23 | 0.00 | **4.88** |
| **SMU_635** | | conserved hypothetical protein | -6.15 | -71.05 | 0.00 | **4.46** |
| **SMU_29** | | phosphoribosylaminoimidazole-succinocarboxamide synthase | -6.52 | -91.97 | 0.00 | **6.96** |
| **SMU_37** | *purH* | phosphoribosylaminoimidazolecarboxamide formyltransferase / IMP cyclohydrolase | -7.42 | -171.20 | 0.00 | **8.61** |

| | | | | | | |
|---|---|---|---|---|---|---|
| **SMU_51** | *purK* | phosphoribosylaminoimidazole carboxylase, ATPase subunit | -9.08 | -542.49 | 0.00 | **5.79** |

**Supplemental Table 3.** ***Strains used in this study*.**

| Species | Strain | Description | Antibiotic resistance | Reference |
|---|---|---|---|---|
| ***S. mutans*** | UA159 | Wild-type *S. mutans* and isogenic parent strain of mutants used in this study | None | Lab Stock |
| | Δ*alsS*Δ*aldB* | Allelic exchange of SMU.1451 and SMU.1452 with *ermB* providing resistance to erythromycin | Erythromycin | This study |
| | Δ*adhC* | Allelic exchange of SMU.129 with *ermB* providing resistance to erythromycin | Erythromycin | This study |
| | Δ*lplA* | Allelic exchange of SMU.131with *ermB* providing resistance to erythromycin | Erythromycin | This study |
| | ΔSMU.1155 | Allelic exchange of SMU.1155, which has no measurable transcptional changes in CDM-Saliva with *ermB* providing resistance to erythromycin (control strain expected to show no fitness changes from the parental) | Erythromycin | (1) |
| | Δ*mntH* | Allelic exchange of SMU.770c with *ermB* providing resistance to erythromycin | Erythromycin | This study |
| | Δ*ccpA* | Allelic exchange of SMU.1591 with *ermB* providing resistance to erythromycin | Erythromycin | This study |
| | Δ*purD* | Allelic exchange of SMU.48 with *ermB* providing | Erythromycin | This study |

| | | | | |
|---|---|---|---|---|
| | | resistance to erythromycin | | |
| | ΔSMU.677 | Allelic exchange of SMU.677 with *ermB* providing resistance to erythromycin | Erythromycin | This study |
| | ΔSMU.678 | Allelic exchange of SMU.678 with *ermB* providing resistance to erythromycin | Erythromycin | This study |
| | ΔSMU.678Δ679 | Allelic exchange of SMU.678 and SMU.678 with *ermB* providing resistance to erythromycin | Erythromycin | This study |
| | UA159 pPGS | Plasmid pPGS integrated into the chromosome carrying aad9 providing resistance to spectinomycin for utilization in coculture CFU experiments | Spectinomycin | Lab Stock |
| | SMU098 | SM1 | None | (2, 3) |
| | SMU101 | U2B | None | (2, 3) |
| | SMU104 | SA41 | None | (2, 3) |
| | SMU105 | SF12 | None | (2, 3) |
| | SMU107 | R221 | None | (2, 3) |
| | SMU108 | M230 | None | (2, 3) |
| | SMU109 | OMZ175 | None | (2, 3) |
| ***S. gordonii*** | DL1 | | None | Lab Stock |
| ***S. sanguinis*** | SK36 | | None | Lab Stock |

**Supplemental Table 4. *Primers used in this study*.**

Underline designates BamHI restriction site, used in PCR ligation mutagenesis approach .

| Primer Name | Primer Sequence (5'-3') | Tm°C |
|---|---|---|
| **△alsSaldB_A** | GGC GAA GCA GAA TTC TGG | 60.5 |
| **△alsSaldB_B** | GAG GAT CCT GAT CAT ACT TTT GGT | 62.2 |
| **△alsSaldB_C** | GAG GAT CCT GAT CAT ACT TTT GGT | 59 |
| **△alsSaldB_D** | CAA GAG CCC TTG GTC ATC | 60 |
| **SMU.678_A** | TGG AGC TTA TCT AAG CTC ATG AGG | 56 |
| **SMU.678_B** | CTC GGA TCC TTC CAG ATC AGG CAC | 61 |
| **SMU.678_C** | TTG GGA TCC GAT GAT ATG GCA GAC | 59 |
| **SMU.678_D** | CGA AAT CAC CAA GTC TAT CAC GCC | 58 |
| **SMU.677_A** | GCG CTT TTG GTA TTG CTG AGC AGC | 61 |
| **SMU.677_B** | GAA GGA TCC CCG CTG ATA TGT CAG | 59 |
| **SMU.677_C** | AGA GGA TCC TCA CAG AAA GTG AGG | 58 |
| **SMU.677_D** | AAA TGT CTG CCA TAT CAT CGT CGG | 57 |
| **△mntH_A** | CGA AGA GCA AGA TCC AAA CGA GTC | 58 |
| **△mntH_B** | TGA GGA TCC TCG CTG AGT GAT TGC | 61 |
| **△mntH_C** | CTG GGA TCC CAA TCC TCA ACT TGC | 60 |
| **△mntH_D** | GGA CTT AGT CCT GTA CAG TAC AGA ACG | 58 |
| **△purD_A** | TTG AAT CTC TGC CAC ATA GAC GAG | 56 |
| **△purD_B** | GGC GGA TCC ATG CTC ACG ACC ACC | 66 |
| **△purD_C** | TCG GGA TCC CAT CGG AAG CAA GGC | 65 |
| **△purD_D** | TCC TGC TTT GTA GGC ATT AGC AGG | 59 |
| **△ccpA_A** | GTT GGC CAA CAG CAA CAG TTC GTG | 61 |
| **△ccpA_B** | ATC GGA TCC CAT CTT GTA TCA TCC | 56 |
| **△ccpA_C** | TGA GGA TCC AGC AGT TGC TAT GCG | 61 |
| **△ccpA_D** | TCG CAA GTG CTC CTG TAC TTC AGG | 61 |
| **SMU_129_A** | CGC TCA AAC TGG ACT TCG TCC | 58 |
| **SMU_129_B** | TTC GGA TCC CTG CAT ATC AAC ACC | 59 |
| **SMU_129_C** | CTT GGA TCC TGG CAT GAA CGG CGC | 65 |
| **SMU_129_D** | AGT TCA ATG CCG ACA ACA CCA CCG | 62 |
| **SMU_131_A** | CTG CTA TGG ACA AGG AAG TAT CGC | 57 |

| | | |
|---|---|---|
| **SMU_131_B** | TTC GGA TCC TAT ATT AAA AGC TGG | 53 |
| **SMU_131_C** | AAG GGA TCC CCA AAG CGA TTG TGG | 62 |
| **SMU_131_D** | GAA ACC GTA CGT GAG ACA ATC TGC | 58 |

**Supplemental Table 5.** ***RNA-seq read counts and mapping statistics*** of reads assigned to *Streptococcus mutans* UA159.

| Sample # | Sample Name | Total Reads / Per Sample | Total Reads Mapped to UA159 Genome using Bowtie2 | Alignment Rate |
|---|---|---|---|---|
| 1 | CDM_1 | 42107998 | 41022837 | 97% |
| 2 | CDM_2 | 26464306 | 25101289 | 95% |
| 3 | CDM_3 | 28922358 | 28080643 | 97% |
| 4 | CDM_Saliva_1 | 28981196 | 27440573 | 95% |
| 5 | CDM_Saliva_2 | 29735504 | 28687977 | 96% |
| 6 | CDM_Saliva_3 | 26209286 | 22229600 | 85% |

**Supplemental Table 6.** ***Tn-Seq read processing and alignment statistics*** for each sample, including initial reads assessed by FastQC, reads entering and retained after Mag6 trimming, reads mapped to the *Streptococcus mutans* UA159 genome using Bowtie2, and reads assigned to genes using HTSeq. Mag6 retention percentages are calculated relative to reads entering Mag6 trimming; alignment and gene assignment percentages are calculated relative to retained reads.

| Sample | Pretrim reads[1] | Reads entering Mag6 trimming | Reads retained after Mag6 trimming | Mag6 retention (%)[2] | Reads mapped to UA159[3] | Alignment rate (%)[4] | Reads assigned to genes[5] | Gene assignment rate (%)[4] |
|---|---|---|---|---|---|---|---|---|
| **BHI_1** | 106,770,631 | 106,303,212 | 99,890,419 | 93.97% | 99,291,754 | 99.40% | 75,241,455 | 75.32% |
| **BHI_2** | 72,390,936 | 72,074,041 | 65,018,840 | 90.21% | 64,444,913 | 99.12% | 47,906,810 | 73.68% |
| **BHI_3** | 96,989,745 | 96,559,625 | 89,821,525 | 93.02% | 88,866,852 | 98.94% | 66,555,477 | 74.10% |
| **CDM_p1_1** | 94,264,488 | 93,848,135 | 86,189,742 | 91.84% | 85,698,267 | 99.43% | 66,741,577 | 77.44% |
| **CDM_p1_2** | 81,640,528 | 81,279,706 | 76,853,899 | 94.55% | 76,490,444 | 99.53% | 60,000,723 | 78.07% |
| **CDM_p1_3** | 68,073,594 | 67,776,329 | 63,721,299 | 94.02% | 63,365,930 | 99.44% | 47,062,641 | 73.86% |
| **CDM_p5_1** | 620,450 | 617,642 | 574,334 | 92.99% | 572,133 | 99.62% | 480,637 | 83.69% |
| **CDM_p5_2** | 132,994,044 | 132,413,304 | 123,610,241 | 93.35% | 122,514,426 | 99.11% | 88,661,326 | 71.73% |
| **CDM_p5_3** | 121,053,351 | 120,523,664 | 114,164,191 | 94.72% | 113,315,641 | 99.26% | 83,497,854 | 73.14% |
| **CDM_HSp1_1** | 6,175,434 | 6,148,622 | 5,736,662 | 93.30% | 5,711,031 | 99.55% | 4,644,708 | 80.97% |
| **CDM_HSp1_2** | 105,127,366 | 104,662,599 | 96,877,582 | 92.56% | 96,230,887 | 99.33% | 71,946,796 | 74.27% |
| **CDM_HSp1_3** | 9,884,714 | 9,841,074 | 9,197,801 | 93.46% | 9,157,568 | 99.56% | 7,378,135 | 80.22% |
| **CDM_HSp5_1** | 52,630,811 | 52,395,754 | 49,424,990 | 94.33% | 49,243,882 | 99.63% | 39,516,434 | 79.95% |

| | | | | | | | | |
|---|---|---|---|---|---|---|---|---|
| **CDM_HSp5_2** | 90,878,918 | 90,481,560 | 85,324,537 | 94.30% | 84,859,663 | 99.46% | 65,014,860 | 76.20% |
| **CDM_HSp5_3** | 2,867,551 | 2,854,410 | 591,539 | 20.72% | 587,497 | 99.32% | 433,078 | 73.21% |

[1] Pretrim reads are the per-sample sequences reported by FastQC after FASTQ Groomer
[2] Mag6 retention = reads retained after Mag6 trimming ÷ reads entering Mag6 trimming × 100.
[3] Mapped reads = reads with exactly one alignment + reads with more than one alignment.
[4] Alignment rate and gene assignment rate both use reads retained after Mag6 trimming (the Bowtie2 input) as the denominator.
[5] Gene-assigned counts are taken from the HTSeq summary __aligned category.
